# Human Lung Alveolar Model with an Autologous Innate and Adaptive Immune Compartment

**DOI:** 10.1101/2025.02.27.640440

**Authors:** Linda Steinacher, Bruno Gjeta, Marisa Pimentel Mendes, Floriana Cremasco, Irineja Cubela, Marina Bellavista, Laura Gaspa Toneu, Luisa Lauer, Qianhui Yu, Ryo Okuda, Barzin Y. Nabet, Velimir Gayevskiy, Michel Siegel, Axel Ducret, Evodie Lassalle, Giacomo Lazzaroni, Jonas Nikoloff, Miguel Camacho Rufino, Christelle Zundel, Leo Kunz, Tamara Zimmermann, Bilgenaz Stoll, Cyrill Roth, Adrian B. Roth, Rajat Mohindra, Nadine Stokar-Regenscheit, Nikolche Gjorevski, Armin Braun, Timothy Recaldin, J. Gray Camp, Lauriane Cabon

**Author notes:** Correspondence to J. Gray Camp and Lauriane Cabon. These authors contributed equally (Linda Steinacher, Bruno Gjeta).

## Abstract

Lung-resident immune cells, spanning both innate and adaptive compartments, preserve the integrity of the respiratory barrier, but become pathogenic if dysregulated^1^. Current in vitro organoid models aim to replicate interactions between the alveolar epithelium and immune cells but have not yet incorporated lung-specific immune cells critical for tissue residency^2^. Here we address this shortcoming by describing human lung alveolar immuno-organoids (LIO) that contain an autologous tissue-resident lymphoid compartment, primarily composed of tissue-resident memory T cells (TRMs). Additionally, we introduce lung alveolar immuno-organoids with myeloid cells (LIOM), which include both TRMs and a macrophage-rich alveolar myeloid compartment. The resident immune cells formed a stable immune-epithelial system, frequently interacting with the epithelium and promoting a regenerative alveolar transcriptomic profile. To understand how dysregulated inflammation perturbed the respiratory barrier, we simulated T-cell-mediated inflammation in LIOs and LIOMs and used single-cell transcriptomic analyses to uncover the molecular mechanisms driving immune responses. The presence of innate cells induced a shift in T cell identity from cytotoxic to immunosuppressive, reducing epithelial cell killing and inflammation. Based on insights obtained with bulk RNA-seq data from the phase 3 IMpower150 trial, we tested whether LIOM cultures could model clinically-relevant but poorly understood pulmonary side effects caused by immunotherapies such as the checkpoint inhibitor atezolizumab^3^. We observed a decrease in immunosuppressive T cells and identified gene signatures that matched the transcriptomic profile of patients with drug-induced pneumonitis. Given its effectiveness in capturing outcomes and mechanisms associated with a prevalent pulmonary disease, this system unlocks opportunities for studying a wide range of immune-related pathologies in the lung.

## Introduction

Organoids have revolutionised our understanding of epithelial biology, providing unique insight into the function and behaviour of epithelial cells in health and disease^4^. However, the breadth of research questions that organoids can be used for is inherently limited by the epithelial exclusivity of the culture composition and the lack of key, closely-associated mesenchymal components^5^. In mucosal barrier tissues like the lung, resident innate and adaptive immune cells perform fundamental functions as homeostatic sensors, orchestrating downstream responses upon detection of epithelial stress^6,7^. Such interactions are critical for maintaining tissue integrity, and their immune-epithelial perturbations are associated with pathological damage and autoimmunity^8^. Accordingly, there is a great need to broaden the applicability of organoid systems and incorporate diverse tissue-specific immune cells in order to facilitate deep interrogation of critical physiological and pathological biological processes^9^.

In vitro lung alveolar models can be derived from human adult or induced pluripotent stem cells^10–12^. The incorporation of blood-derived immune cells such as peripheral blood mononuclear cells (PBMCs)^13–15^ or macrophage cell lines like THP-1^16–18^ has been achieved but the relevance of immortalised or blood-derived immune cells in this context is unclear. In the alveoli, cuboidal alveolar epithelial type 2 (AT2) cells produce surfactant and act as alveolar stem cells^19^ that proliferate and differentiate into squamous alveolar epithelial type 1 (AT1) cells, the flat, elongated cells responsible for gas exchange. AT2s are notoriously difficult to maintain in standard 2D cultures^20^, but organoid protocols enable the generation of 3D alveolospheres^10–12^, providing an appropriate baseline for the addition of further relevant cell types. In the alveoli, tissue-resident alveolar macrophages^21^ and tissue-resident memory T cells (TRMs)^22–24^ are of particular interest due to their abundance and critical functions.

We have previously developed and validated human intestinal immune organoids formed by self-assembly of epithelial organoids and autologous TRMs^25^. In this study, we established protocols for the simultaneous isolation of resident myeloid, lymphoid and epithelial AT2 cells from adult human lung tissue and subsequently assembled these cells to generate an immunocompetent in vitro model of the human alveolar region. We used diverse readouts to reveal an unanticipated role for myeloid cells in regulating pathogenic T cell-mediated inflammation, and further applied the model to interrogate clinically-relevant but poorly understood pulmonary side effects induced by targeted cancer therapies and immuno-modulators^3^, namely the EGFR inhibitor erlotinib and the PD-L1 blocker atezolizumab.

## Results

To better understand the interactions between lung epithelium and tissue-resident immune components, we visualised and quantified immune cells present in histological patient-derived sections of normal lung tissue. We noted the integration of T lymphocytes (CD3^+^) and myeloid cells (CD68^+^) in both healthy and inflamed areas, proximal to and closely interacting with the alveolar epithelium (**Fig. 1a** and **Extended Data Fig. 1a, b**). As expected, the number of lymphoid and myeloid cells per area increased with elevated levels of inflammation, suggestive of an active local immune response (**Extended Data Fig. 1c-e**). The AT2 cell-specific marker (HT2-280) increased in inflamed tissue, consistent with the reparative function of these epithelial cells and the physiological hyperplastic reaction when reacting to an inflammatory environment in the alveoli (**Extended Data Fig. 1c**). Guided by these observations (**Fig. 1a)** we sought to recreate the alveolar region in vitro and thus established a workflow to isolate three different cell lineages from the same tissue resection (**Fig. 1b**): AT2 cells for the generation of alveolar organoids^10^, tissue-resident memory T cells (TRMs) isolated using a scaffold-based crawl-out protocol^25,26^, and alveolar myeloid cells (AMs).

**Figure 1.**
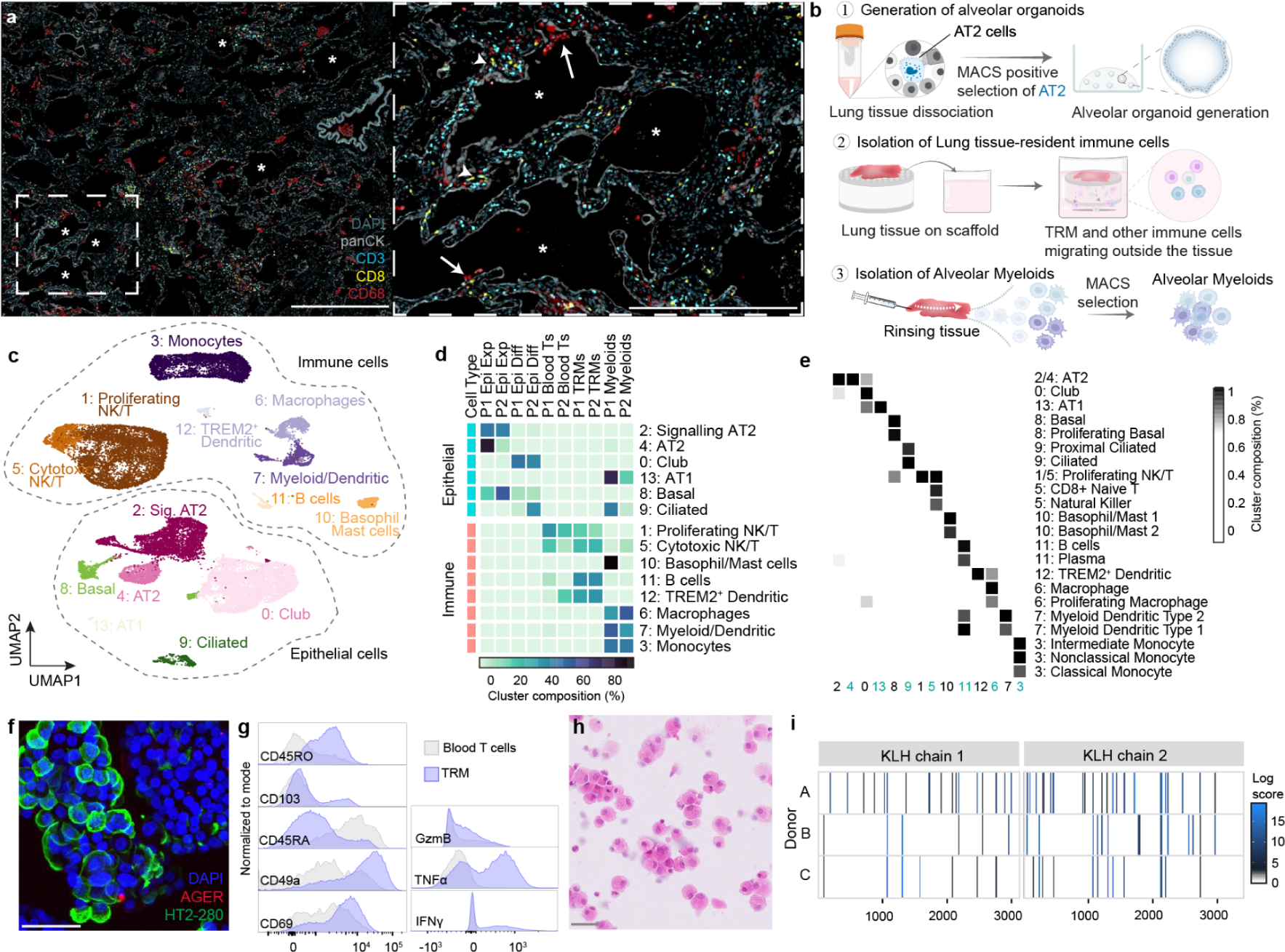
Autologous workflow demonstrates the feasibility of isolating different cell types from fresh lung tissue resections. **a**, Representative multiplex immunofluorescence (IF) image of patient-derived normal adjacent to tumour lung tissue showing epithelial cells (panCK - grey) and immune cells such as T cells (CD3 - cyan, CD8 - yellow) and myeloid cells (CD68 - red). Stars indicate alveoli, arrows indicate accumulations of myeloid cell clusters and arrowheads show T cell clusters. Scale bars, left 1 mm, right 500 µm. **b**, Illustration of the autologous workflow to isolate the 3 different cell types. **c,** UMAP embedding of single-cell transcriptional profiles of isolated cells coloured by cluster and cell type. **d**, Heatmap of the proportional distribution of clusters across all 10 sequenced samples. 2 donors per sample type. **e,** Heatmap of transcriptional signatures from a public lung cell atlas (Travaglini et al.^27^) (y-axis) on clusters identified in **c** (x-axis). **f**, Immunofluorescent images of alveolar organoids in the expansion medium, showing retention of alveolar epithelial type 2 cells (HT2-280 - green) and no upregulation of AGER (red). Scale bars, 50 µm. **g**, Representative flow cytometry histograms comparing TRMs (blue) with donor-matched blood-derived T cells (grey), demonstrating higher expression of tissue-resident markers (CD69, CD49a, CD103) and memory marker (CD45RO) on TRMs at baseline. Upon stimulation, TRMs exhibit higher intracellular cytokine upregulation (GzmB, TNFα, IFNγ). Gated on viable, CD45+, CD3 single cells. **h**, Haematoxylin and eosin staining of a cytospin of alveolar myeloid extracts shows clusters of large, 30-40 µm round cells with eosinophilic granular to foamy cytoplasm and round to bean-shaped nuclei. Scale bar, 50 µm. **i**, Heatmap depicting the cluster profile of MHC-associated peptide proteomics identified peptides for KLH in 3 different donors. Sequence regions are organised according to the protein domains. Identified peptide clusters are shown as coloured regions with varying abundances per sequence position (log scale).

We used single-cell RNA-sequencing (scRNA-seq) to analyze different alveolar fractions (expanded and differentiated epithelial, lymphoid, myeloid) from two independent donors, and included donor-matched blood T cells as non-lung immune lymphoid cell comparison. Heterogeneity analysis identified 14 molecularly distinct clusters (**Fig. 1c** and **Extended Data Fig. 2a**), with both donors contributing to each cluster, indicating low donor-to-donor variability (**Fig. 1d** and **Extended Data Fig. 2b**). Clusters were annotated using a human lung cell atlas as a reference comparison^27^, revealing epithelial cells originating from the expanded alveolar organoid sample were predominantly AT2 cells with a minor contribution of basal cells (**Fig. 1e**). Organoids from the same donor grown in differentiation conditions, contained a low abundance of AT1 and AT2 cells, and were enriched in club cells and, to a lesser extent, to ciliated and basal cells (**Fig. 1e**). TRMs and matched blood-derived T-cell samples were composed of similar cell types. The alveolar myeloid cells consisted predominantly of monocytes (intermediate, non-classical, classical), DCs and macrophages, with a small fraction of epithelial cells (2-10% depending on the donor) obtained during the tissue lavage.

Visualisation of the most highly enriched genes per cluster supported cell-specific activities (**Extended Data Fig. 2c**). Expanded alveolar organoids (c2 and c4) expressed AT2-related genes such as *SFTPA2*, *SFTPA1*, *SFTPB*, *SFTPC*, indicative of surfactant production. Differentiated alveolar organoids conversely expressed club cell-like genes such as *SCGB3A2,* necessary for the immune regulation of the upper airways. Alveolar myeloid cells expressed genes pertinent for antigen presentation (*HLA-DP, HLA-DQ, HLA-DR*) inflammation and phagocytosis (*PPARG, MRC1* and *SIGLEC1*) (**Extended Data Fig. 2d**). As the TRMs and blood T cells showed a certain degree of gene overlap, we performed a neighbourhood analysis to identify more subtle differences between these two populations (**Extended Data Fig. 2e,f**). Blood-derived T cell neighbourhoods, showed enriched expression of central memory T cells genes *ID3*, *CCR7*, *TCF7* and *SELL* (**Extended Data Fig. 2g**), whereas the predominant TRM neighbourhood expressed genes related to tissue residency (*ITGA4, ITGA2, ITGA1*) or adhesion (*LGALS3*), which were lowly expressed by the blood-derived T cells (**Extended Data Fig. 2h**).

We stained the expanded and differentiated organoids by immunofluorescence and found that the expanded organoids retained AT2 cells (HT2-280^+^), as expected from transcriptional analysis (**Fig. 1f**). Since AT2 cells possess innate immune functions, including antigen presentation and cytokine production^28,29^, we chose to grow organoids in the expansion medium for all downstream applications.

Flow cytometric analysis showed that TRMs greatly differed from donor-matched blood-derived T cells (**Fig. 1g**) through expression of surface molecules associated with tissue retention (CD69), extracellular matrix association (CD49a, encoded by *ITGA1*) and epithelial cell integration (CD103, encoded by *ITGAE*), as well as the short isoform of CD45 expressed on antigen-experienced memory lymphocytes (CD45RO). Blood T cells however, presented higher expression of the long isoform of CD45 (CD45RA), usually expressed by naive T cells, and less or none of the residency markers (**Fig. 1g** and **Extended Data Fig. 3a**). Since TRMs are reported to rapidly produce effector molecules in response to appropriate cues, we assessed the cytokine production capacity of our isolated TRMs following stimulation. As anticipated, we observed increased expression of granzyme B, TNFα and IFNγ compared to their matched blood counterparts (**Fig. 1g** and **Extended Data Fig. 3b**).

Haematoxylin and eosin (H&E) assessment of our isolated myeloid compartment revealed cells of approximately 30 to 40 µm in size with granular to foamy cytoplasms (**Fig. 1h**). Incubating the cells with the foreign keyhole limpet hemocyanin (KLH) protein demonstrated their capacity to phagocytose, cleave and subsequently present KLH peptides via HLA class II complexes (**Fig. 1i**). The AMs were equally capable of engulfing *Staphylococcus aureus* bioparticles and produced vast quantities of various pro-inflammatory factors at baseline and in response to stimulation (**Extended Data Fig. 3c-f**). Accordingly, the isolated myeloid and lymphoid compartment retained their functional activity upon removal from the tissue, and demonstrated that inclusion of resident immune cells was an essential pre-requisite for our model, in order to truly capture organ-specific inflammatory responses.

Having established an autologous workflow that allowed for the isolation of the different cell types, which retained tissue-specific functionality following removal from the tissue, we combined these three elements to form an immune-competent alveolar organoid system. Imaging of the cultures after establishment revealed extensive homeostatic interactions between the different components (**Fig. 2a** and **Extended Data Fig. 4a**), with T cells regularly adopting asymmetrical elongated shapes to contact the epithelium. TRMs also exhibited a dynamic behavior in the culture, migrating between the ECM and multiple alveolar organoids (**Supplementary Video 1**). AMs remained mainly in the extracellular matrix, but occasionally interacted with the organoid surface (**Fig. 2b**). Quantification demonstrated that CD4^+^ and CD8^+^ TRMs interacted equally with the organoids, representing about 8% of the total lymphoid population. By comparison, 2% of the AMs interacted with epithelial cells (**Fig. 2c**). T cells in close contact with the epithelium often expressed the E-cadherin receptor CD103, likely supporting their interaction and integration with the alveolar surface (**Fig. 2d** and **Extended Data Fig. 4c, d**).

**Figure 2.**
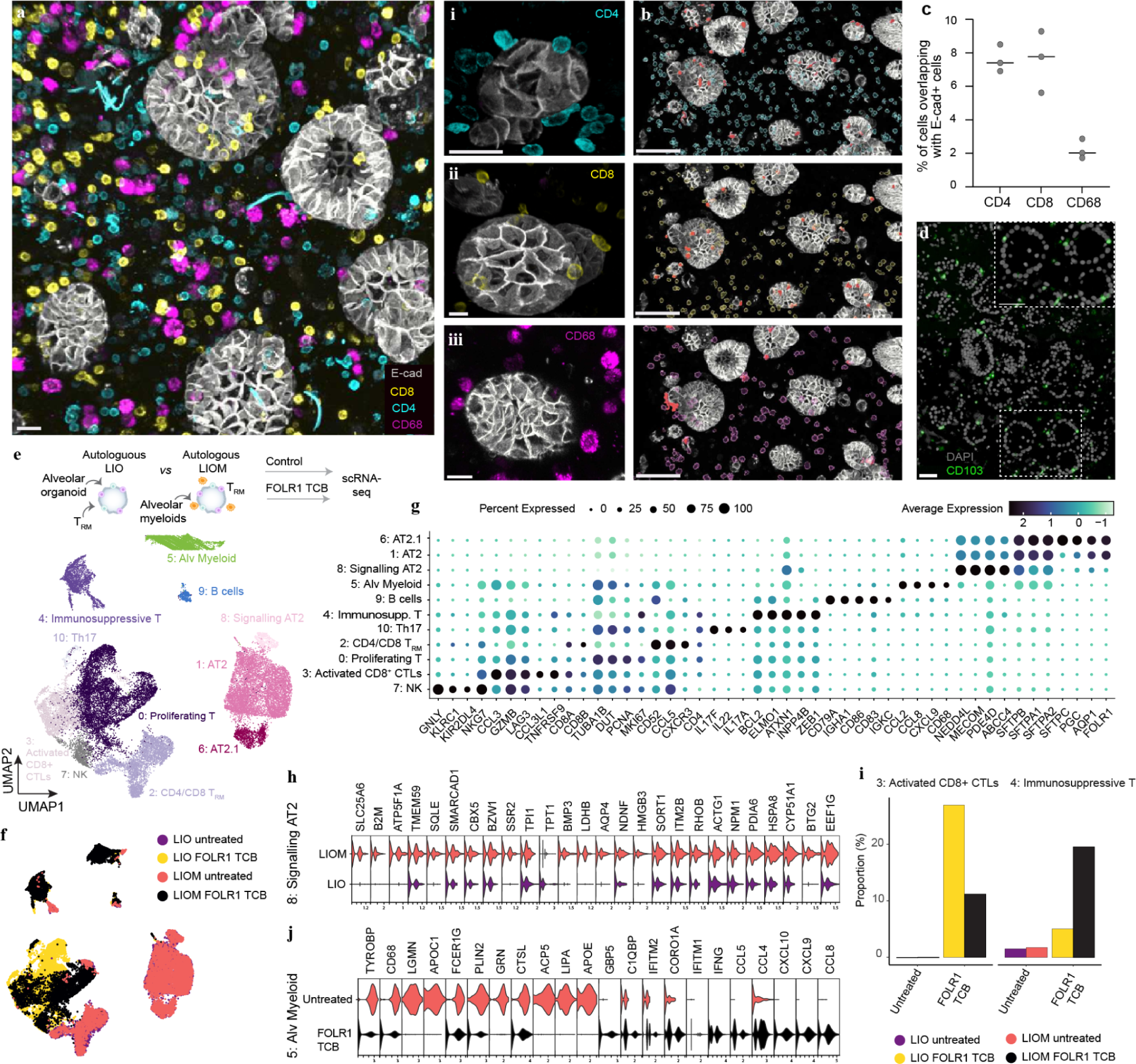
Human lung alveolar model with an autologous innate and adaptive immune compartment is characterized at baseline and upon T cell-mediated inflammation. **a**, Representative maximum projection immunofluorescence (IF) image of a fixed tri-culture after 72 h of culture. Organoids in white (E-cad), CD8 in yellow, CD4 in cyan and myeloid cells (CD68) in magenta. Close-up of CD4 (i), CD8 (ii) and CD68 (iii). Scale bars, 20 µm. **b**, Maximum projection IF panels of immune cells interacting with E-cad^+^ cells (red). Top CD4 (i), middle CD8 (ii), and bottom CD68 (iii). Scale bars, 100 µm. **c**, Quantifications of **a** and **b** showing the percentage of immune cells interacting with the epithelium. Line at median, technical triplicates. **d**, Multiplex IF staining of an FFPE tri-culture after 72 h of culture demonstrating the interaction of CD103^+^ immune cells (green) with organoids (white). Scale bar, 50 µm. **e**, Integrated UMAP plot of scRNA-seq data from 4 samples as shown in the illustration (LIO and LIOM at baseline and after FOLR1 TCB stimulation at 72 h) from one donor colored by 11 clusters (**e**) or by sample (**f**). **g**, Dotplot summarising the expression patterns of representative genes across clusters identified in **e**. **h**, Violin plots showing differences in gene expression of cluster 8 (signalling AT2, top plot) when cultured in the absence (LIO) or presence (LIOM) of myeloid cells. **i**, Barplots displaying the proportional differences of activated CD8^+^ CTL (cluster 3) and immunosuppressive T (cluster 4) in the absence and presence of alveolar myeloid cells with and without treatment. FOLR1 TCB was used at 1 ng/mL. **j**, Violin plots showing differences in gene expression of cluster 5 (alveolar myeloid) with and without FOLR1 treatment.

We performed scRNA-seq to characterise myeloid-lymphoid-epithelial crosstalk within the model’s homeostatic baseline. Specifically, we assessed TRMs co-cultured with alveolar organoids in the presence or absence of AMs, from herein referred to as **l**ung **i**mmuno-**o**rganoids with **m**yeloid cells (LIOM) versus **l**ung **i**mmuno-**o**rganoids (LIO). We additionally included conditions to direct T cell inflammatory activity towards the epithelium by treating LIOM and LIO cultures with a T cell-bispecific antibody (TCB) targeting folate receptor 1 (FOLR1). This TCB was originally envisioned for the treatment of FOLR1-expressing ovarian carcinoma^13,30^. However, on-target off-tumour toxicities prevented its development due to the expression of the FOLR1 antigen by healthy epithelial cells, such as AT2s (**Extended Data Fig. 4e**). Using the FOLR1 TCB to induce T cell-mediated epithelial cell damage allowed us to observe how the interactions between the different cell lineages change during the course of inflammation.

The sequencing data from the different conditions elucidated 11 distinct clusters that could be assigned to myeloid (c5: alveolar myeloid), lymphoid (c0: proliferating T cells, c2: CD4/CD8 TRMs, c3: activated CD8 cytotoxic T lymphocytes (CTLs), c4: immunosuppressive T cells, c7: NK cells, c9: B cells, c10: Th17 cells) and epithelial cell types (c1: AT2, c6: AT2.1, c8: signalling AT2) (**Fig. 2e-g**). Direct comparison of the baseline LIOM and LIO samples revealed a specific impact of myeloid cells on the signalling AT2 cluster (c8), with altered expression of genes involved in proliferation (*EE1FG, BTG2, LDHB, SLC25A6*), stemness (*HMGB3, BMP3)*, lipid metabolism important for surfactant production (*CYP51A1, SQLE*), intracellular trafficking (*HSPA8, SORT1*) and antigen presentation (*B2M*) (**Fig. 2h**). This suggested that AMs might play a role in modulating the functional and regenerative properties of AT2 cells by altering their gene expression profile across several AT2 specific critical biological processes.

Immune cells in the LIO and LIOMs at baseline had a complete overlap of the gene signatures, suggesting that AMs did not affect TRM identity (**Extended Data Fig. 4f, g**). In contrast, AM cells within the LIOM model drastically altered drug-induced T-cell states. The presence of AMs not only prevented the emergence of cytotoxic TRMs (c3) upon TCB treatment, but also potently promoted the emergence of immunosuppressive TRMs (c4), highlighting a critical shift in T-cell behavior (**Fig. 2i** and **Extended Data Fig. 4h**). Interestingly, alveolar myeloid cells (c5) downregulated markers associated with immune resolution (*APOE, LIPA, GRN, APOC1*), phagocytosis (*CTSL*), bacterial clearance (*ACP5*) and universal macrophage markers (*FCER1G*, *TYROBP* and *CD68*), while simultaneously upregulating IFNγ-induced genes characteristic of a Th1 response (*CXCL9, CXCL10, CCL5*) (**Fig. 2j**).

To investigate the underlying intercellular signaling mechanisms behind those changes, we performed a receptor-ligand network analysis, which revealed a significant increase in putative cellular crosstalk in LIOM cultures compared to LIO cultures (**Fig. 3a**). Gene ontology analysis identified pathways associated with increased immune cell activation in LIO (**Fig. 3a left**), whereas pathways related to the regulation of the Th1 immune response and chemotaxis were upregulated in LIOM (**Fig. 3a right**). The network analysis proposed that alveolar macrophage ligands (c5, module 2) such as CXCL9, CXCL10, and CCL2 signaled to proliferating T cells (c0, c10) and B cells (c9) via CXCR3, and to immunosuppressive T cells (c4) via CCR4, with further interactions via the LGALS3-ITGB1 and ALCAM/CD6 axis. Simultaneously, immunosuppressive T cells (c4) signaled to NK cells, proliferating T cells, cytotoxic T cells, Th17 cells, and B cells via TGFβ, TIGIT pathway, TNF receptors, IL35, and GPR132 (**Fig. 3b, c** and **Extended Data Fig. 5a-c**). These receptor-ligand interactions suggest both direct and indirect effects of AMs on TRMs. Consistent with this, we observed direct cell-to-cell interactions and synapse formation between AMs and TRMs (**Fig. 3d**). Finally, functional validation revealed that this immunosuppressive reprogramming of TRMs was reflected by decreased epithelial apoptosis and a reduction in pro-inflammatory cytokines secretion (IFNγ, IL-6, and granzyme B) in LIOM cultures (**Fig. 3e-g**), confirming the functional impact of AM-mediated modulation of T-cell states.

**Figure 3.**
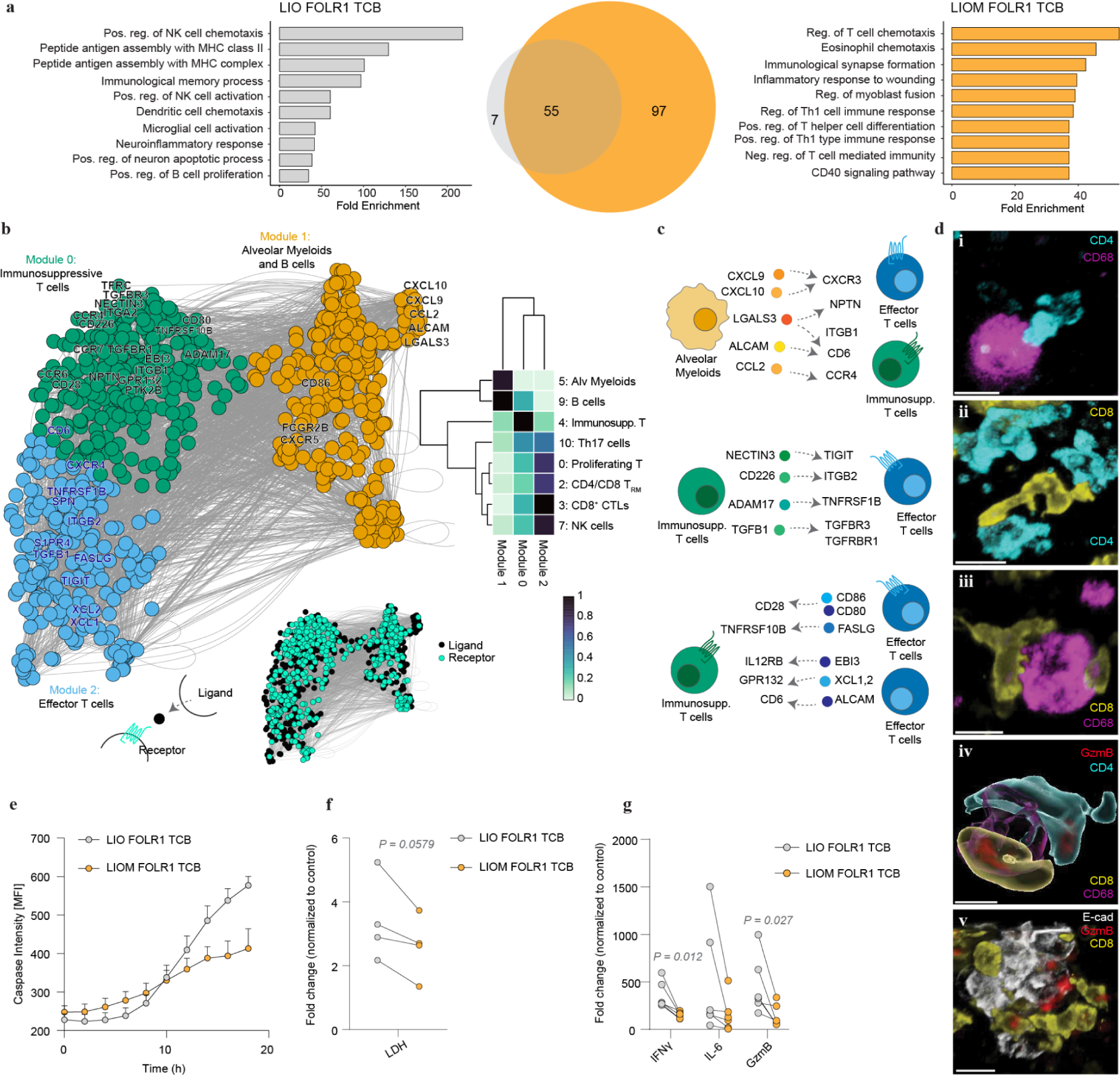
Alveolar myeloid cells dampen TRM-mediated alveolar epithelial damage. **a**, Left, ontology analysis of genes upregulated in FOLR1 TCB-treated LIOs presenting positive upregulation of pro-inflammatory pathways. Middle, Venn diagram showing the unique and overlapping signalling molecules in the LIOs (grey) vs LIOMs (yellow). Right, ontology analysis of genes upregulated in treated LIOMs reveals Th1-related pathways. **b**, Ligand-receptor pairing analysis from scRNA-seq data of the immune cell populations in the LIOMs. Co-expression modules are used to colour ligands and receptors. Representative genes are highlighted. Heatmap (right) shows the average expression of each gene within a module across each cell cluster. **c,** Illustration of selected ligand-receptor interactions. Signaling cells entail alveolar myeloid cells (top), immunosuppressive T cells (middle) and effector T cells (bottom). **d,** Representative maximum projection immunofluorescence (IF) image of fixed tri-culture after 24 h of FOLR1 TCB treatment showing synapse forming cells. i, CD4 (cyan) with CD68 (magenta); ii, CD8 (yellow) with CD4; iii, CD8 with CD68; iv, CD68, CD4 and CD8 cells containing GzmB (red); v, GzmB containing CD8^+^ cells interacting with organoids (E-cad, white). Scale bars, 15 µm. **e**, Bihourly quantification of caspase 3/7 intensity identified within organoids over 20 h in FOLR1 TCB treated LIOs and LIOMs. 3 independent biological replicates, mean and SEM. **f,** Line graphs of the LDH fold change compared to the control in treated LIO and LIOM cultures. Mean of 4 biological independent replicates. Two-tailed paired t-test. **g**, Line graphs showing the fold change of cytokines compared to the control in treated LIO and LIOM cultures. Mean of technical replicates shown for 5 biological independent replicates. Two-tailed paired t-test.

These findings underscore the complex interplay between myeloid and T cells in modulating immune responses within the lung microenvironment. This complexity is particularly relevant in the context of cancer immunotherapy, where immune checkpoint inhibitors, such as those targeting PD-1/PD-L1, have revolutionized the treatment of solid tumors like melanoma and non-small cell lung cancer (NSCLC)^32^. However, these therapies can lead to immune-related adverse events, including checkpoint inhibitor-associated pneumonitis (CIP)^33–35^. A recent meta-analysis found that around 35% of fatalities associated with anti–PD-1/anti–PD-L1 treatments are attributed to CIP^36^. To explore the immune cell dynamics involved in CIP, we analyzed clinical data from the IMpower150 trial, which examined NSCLC patients treated with the anti-PD-L1 checkpoint inhibitor atezolizumab in combination with carboplatin and paclitaxel (ACP). Our analysis of baseline, pre-treatment tumor samples from the ACP arm revealed that patients who developed CIP exhibited higher gene signatures for antigen-presenting cells (APCs), including macrophages and dendritic cells (DCs), as well as elevated MHC class II expression, suggesting that these myeloid subsets may contribute to the development of CIP (**Fig. 4a**)^37–40^. This observation pointed to a potential role for myeloid cells in CIP pathogenesis, making the LIOM model particularly well-suited for further investigation, as it integrates both myeloid cells and T cells in a lung-specific microenvironment.

**Figure 4.**
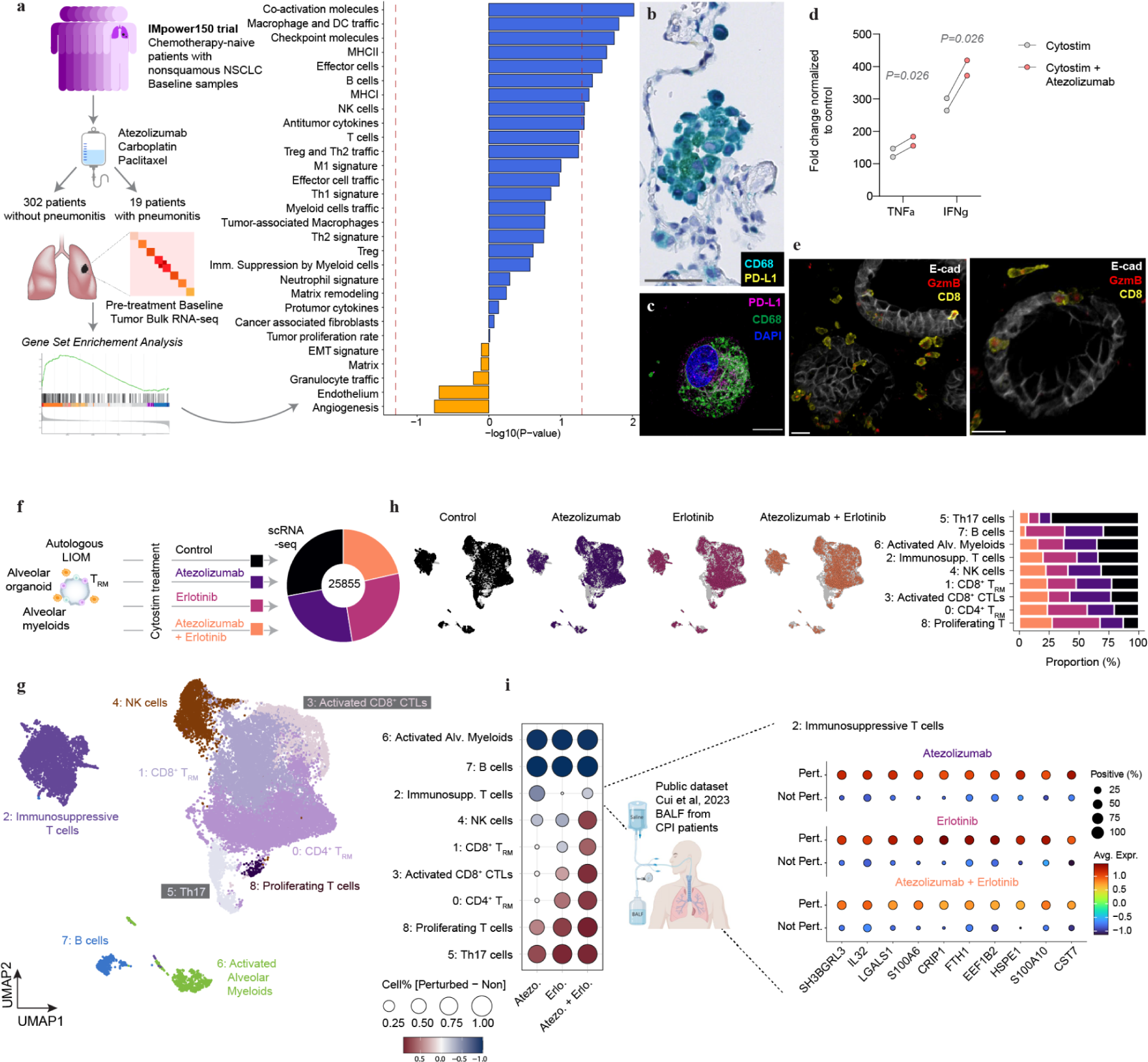
LIOMs can model aspects of checkpoint inhibitor-induced pneumonitis. **a**, Left, illustration of the IMpower150 study and the subsequent inclusion of patients without (302) and with (19) pneumonitis. Right, bulkRNA-sequencing data from baseline lung tumour samples of atezolizumab-treated patients were used to generate the barplot depicting up- and downregulated gene signatures. The red dashed lines indicate P values of 0.05 on a −log_10_ scale. **b**, Representative duplex IHC staining on formalin-fixed paraffin-embedded (FFPE) lung tissue shows CD68^+^ cells (blue) co-expressing PD-L1 (yellow). Scale bar, 50 µm. **c**, Representative IF image of isolated fixed alveolar myeloid cell demonstrates co-expression of CD68 (green) and PD-L1 (magenta). Scale bar, 10 µm. **d**, Line graph presenting the fold change normalized to the control of cytokine expression (IFNγ, TNFα) in CytoStim and CytoStim + atezolizumab treated cultures for 72 h. Mean of technical replicates shown for 2 biological independent replicates. Two-tailed paired t-test. **e**, Two representative maximum projections IF of fixed tri-culture after 72 h of CytoStim + atezolizumab treatment showing CD8^+^ cells (yellow) containing GzmB (red) that are interacting with organoids (white). Scale bars, 20 µm. **f**, Overview of the scRNA-seq experiment of LIOMs with treatments combined with CytoStim (control), resulting in a total of 25855 cells sequenced. **g**, Integrated UMAP plots showing 9 distinct clusters that were annotated based on marker gene expression and comparison to a primary lung cell atlas previously used in Fig. 1. **h**, Left, integrated UMAP shown for each sample separately. Right, stacked barplot shows sample contribution to each cluster as identified in **g**, columns sum to 100. **i**, Left, dotplot showing the perturbed and non-perturbed percentages across all samples and clusters compared to the CytoStim control. Dotsize represented the absolute value of proportion difference between perturbed and unperturbed cells. Colour represents proportion difference. Right, comparing the gene signatures of cluster 2 (immunosuppressive T cells) to a public dataset from Cui et al.^31^, revealing an overlap of signatures found in the treated LIOMs. Dotsize is indicative of percentage of positive cells.

We next used LIOMs to model the immunosuppressive events that could lead to epithelial damage. First, we confirmed the presence of PD-L1 in lung resection tissues by duplex IHC staining and on isolated myeloid cells by immunofluorescence stainings after cryopreservation (**Fig. 4b, c**). Initially, when treated with atezolizumab alone, we did not observe significant increases in pro-inflammatory cytokines compared to controls (**Extended Data Fig. 6a**). However, considering that underlying viral infections have been linked to heightened CIP risk, we modified our experimental approach by applying an MHC-dependent polyclonal stimulus (CytoStim) to promote robust T cell activation. This adjustment led to a marked increase in secreted TNFα and IFNγ following atezolizumab treatment, closely mirroring clinical observations^41^ (**Fig. 4d** and **Extended Data Fig. 6b, c**). We also observed cytotoxic T cells interacting closely with epithelial cells, further validating the inflammatory response seen in the clinical setting (**Fig. 4e**).

Next, we used scRNA-seq to interrogate the transcriptomic dynamics underlying atezolizumab-induced CIP-like outcomes, with a focus on tissue-resident lymphoid and myeloid cells. Clinical reports suggesting that the combination of atezolizumab and the epidermal growth factor receptor (EGFR) tyrosine kinase inhibitor, erlotinib, leads to higher levels of CIP provided the rationale for using this combination. After integrating data from four different samples, we identified 9 distinct cellular clusters (**Fig. 4f, g**). Comparing the effects of three treatments—atezolizumab, erlotinib, and the combination of both—against the CytoStim baseline, we identified differentially enriched genes for each cluster (**Extended Data Fig. 6d**). Notably, Th17 cells (c5), which were predominant in the baseline sample, were significantly reduced with treatment. Immunosuppressive T cells (c2) also showed a smaller, but still notable, decrease. Atezolizumab monotherapy was associated with a higher percentage of CD8^+^ CTLs and NK cells (c3 and c4), while erlotinib maintained a larger proportion of proliferative T cells and CD4^+^ TRM (c8 and c0) (**Fig. 4h**). We assessed the proportion of cells across all clusters that changed in the different samples compared to the CytoStim control, revealing that all clusters were perturbed, with strongest changes in the combination treatment (**Fig. 4i**). To assess the clinical relevance of these changes, we compared the transcriptomic signatures of perturbed cells with published scRNA-seq data from bronchoalveolar lavage samples of CIP patients^36^ (**Fig. 4i**). The signatures found in the immunosuppressive T cell cluster of treated samples overlapped with those from CIP patients **(Fig. 4i)**. Interestingly, the immune response-specific genes *IL-32*, *CRIP1*, and *CST17* were upregulated in all perturbed cells under all treatment conditions, further supporting the idea that certain molecular and cellular mechanisms may be shared by drugs that induce pneumonitis. These findings may also highlight druggable pathways to prevent or mitigate this potentially fatal adverse event.

## Discussion

In this manuscript, we have described an immunocompetent alveolar organoid system composed of human tissue-derived alveolar epithelium and autologous resident lymphocytes with and without myeloid cells. A portion of the resident lymphocytes, mainly CD103^+^ TRMs, integrated into the alveolar epithelium and exerted their cytotoxic and cytokine-releasing functions upon TCR stimulation with a TCB or polyclonal stimulus. This is in line with their function as memory T cells that respond rapidly upon reexposure to lung-specific pathogens^6^. When included, myeloid cells influenced the transcriptome of the epithelium, promoting proliferation and stemness of the AT2 signalling population^27^ in baseline co-culture conditions. The role of myeloid cells in supporting AT2 proliferation and stemness is poorly understood, but has been reported in some mouse models of lung injury and infection^42–44^. Although human Pluripotent Stem Cell (PSC) derived macrophages have been added to PSC derived alveolar organoids, their impact on AT2 biology has not been investigated in that context^45,46^. The AT2 signalling findings we observed in LIOMs versus LIOs may however be an indirect effect of alveolar myeloids via their impact on TRMs. Utilizing an in vitro TRM-AT2 co-culture system^47^, TRMs were indeed found to inhibit AT2 growth, and the phenotype was reversible with addition of neutralizing IFNγ antibody in line with previous findings that interferons suppress AT2 organoid growth^10,48^. Therefore a plausible mechanism of alveolar myeloids in LIOMs could be the dampening of IFNγ signaling in TRMs resulting in improved AT2 maintenance.

Myeloid cells were observed physically interacting with T cells, forming what resembles immune synapses upon TCR stimulation. Ligand-receptor network analysis suggested that these cells were initially recruited to each other via chemokines, particularly the CXCR3 axis, and then maintained via conventional TCR and co-receptor cell-to-cell modules. These interactions did not modulate the TRM transcriptomic profile at baseline but intriguingly exerted a striking effect on their function upon stimulation, limiting epithelial damage upon TCB treatment by reducing IFNγ signalling and inducing immunosuppressive T cells. This effect was likely more of a delay than a blockade as alveolar myeloids started to shift towards a pro-inflammatory phenotype in culture over time. Collectively, our observations positioned AMs as critical resolvers of lung inflammation, limiting T cell-induced epithelial damage and promoting alveolar regeneration.

Detailed analysis of receptor-ligand pairs of alveolar myeloid and lymphoid cells in TCB-treated LIOMs highlighted expected antigen presenting cell-T cell interactions such as CXCL9/10-CXCR3 or CD86/CD80-CD28, but also tissue-resident or even lung-specific pairs, confirming the tissue-resident properties of our in vitro approach. For instance, IL35 is involved in the suppressive function of regulatory T cells and can promote an anti-inflammatory microenvironment within lung tissue^49^. Selective myeloid depletion of *LGALS3* has been reported to protect mice against acute and chronic lung injury^50^, and *LGALS3* has been found to be upregulated in the lungs of patients with idiopathic pulmonary fibrosis^51^, chronic obstructive pulmonary disease (COPD), asthma and viral-induced acute respiratory distress syndrome. *GPR132* has recently been proposed to be a signal within the tissue that controls the quantity and quality of CD8^+^ TRM pools^52,53^. We propose that the use of LIOs and LIOMs may reveal novel tissue-specific immune biology and provide a platform for comprehensive characterisation of such pathways.

Checkpoint-inhibitor associated pneumonitis (CIP) is a potentially fatal side effect of treatment with immunomodulatory agents such as the PD-L1 blocker atezolizumab. Here, we showed that LIOMs treated with checkpoint inhibitors and EGFR tyrosine kinase inhibitors in the context of a polyclonal T cell activation were able to capture certain patient-relevant features, such as those observed in the bronchoalveolar lavage fluid (BALF) of CIP patients^31^. Specifically, we observed a push towards cytotoxic effector T cells and Th1 responses together with a reduction of immunosuppressive T cells under drug treatment conditions. We also found an overlap between the gene signatures of immunosuppressive T cells in LIOM cultures and those of CIP patients. Regulatory T cells have been proposed to contribute to the onset of CIP but both reduction and expansion have been reported^41,54^. Interestingly we observed a decrease of Th17 cells upon treatment which is in apparent contradiction with a study performed in BALF and serum of patients with CIP showing increased IL17A and Th17 cells^54^. Additional experiments would be needed to further understand the molecular mechanisms behind that discrepancy.

We propose that our model could be further exploited to discover biomarkers or treatments to mitigate drug-induced pneumonitis. Indeed, our system presents the advantage of comparing perturbations within the same donor background under all treatment conditions, unlike what is possible in a clinical setting. In sum, by incorporating a human autologous innate and adaptive compartment, we have enhanced alveolar organoids with features critical for studying the dysregulation of respiratory immunity that underlies diseases such as asthma, COPD, respiratory infections and cancer.

---

**Extended Data Figure 1:**
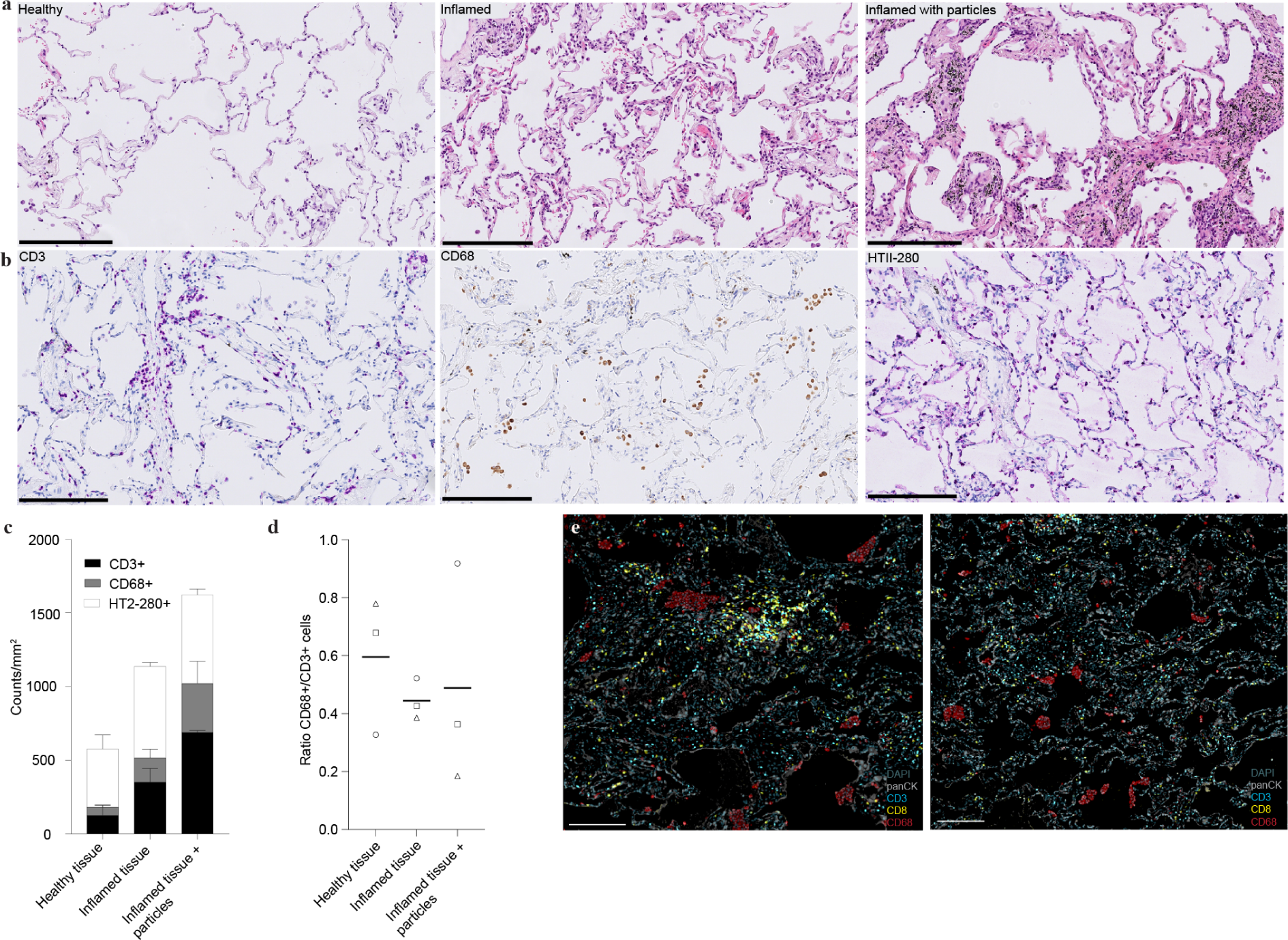
Stainings and quantifications of native lung tissue guides the development of the autologous in vitro model. **a**, Haematoxylin & eosin staining of FFPE lung tissue and subsequent identification of different regions on the same section. Left, healthy region. Middle, inflamed region. Right, inflamed region with black pigments in aggregated macrophages. Scale bars, 250 µm. **b**, IHC staining of sequential sections of FFPE lung tissue with different markers. Tissue stained for CD3 in purple (left), CD68 in brown (middle), and HT2-280 in purple (right). Scale bars, 250 µm. **c**, Quantifications of the sequential IHC stainings from **b** per mm^2^ for the three different regions shown in **a**. Mean and SEM of 3 independent biological replicates shown. **d**, Line plots displaying the mean of the ratios of CD68 to CD3 cells based on the results of **c**. The different symbols indicate the three different donors. **e**, Multiplex immunofluorescence images on FFPE lung tissue showing two inflamed regions with accumulations of CD3 (cyan), CD8 (yellow) (left) and CD68 (red) cells (right). Scale bars, 250 µm.

**Extended Data Figure 2:**
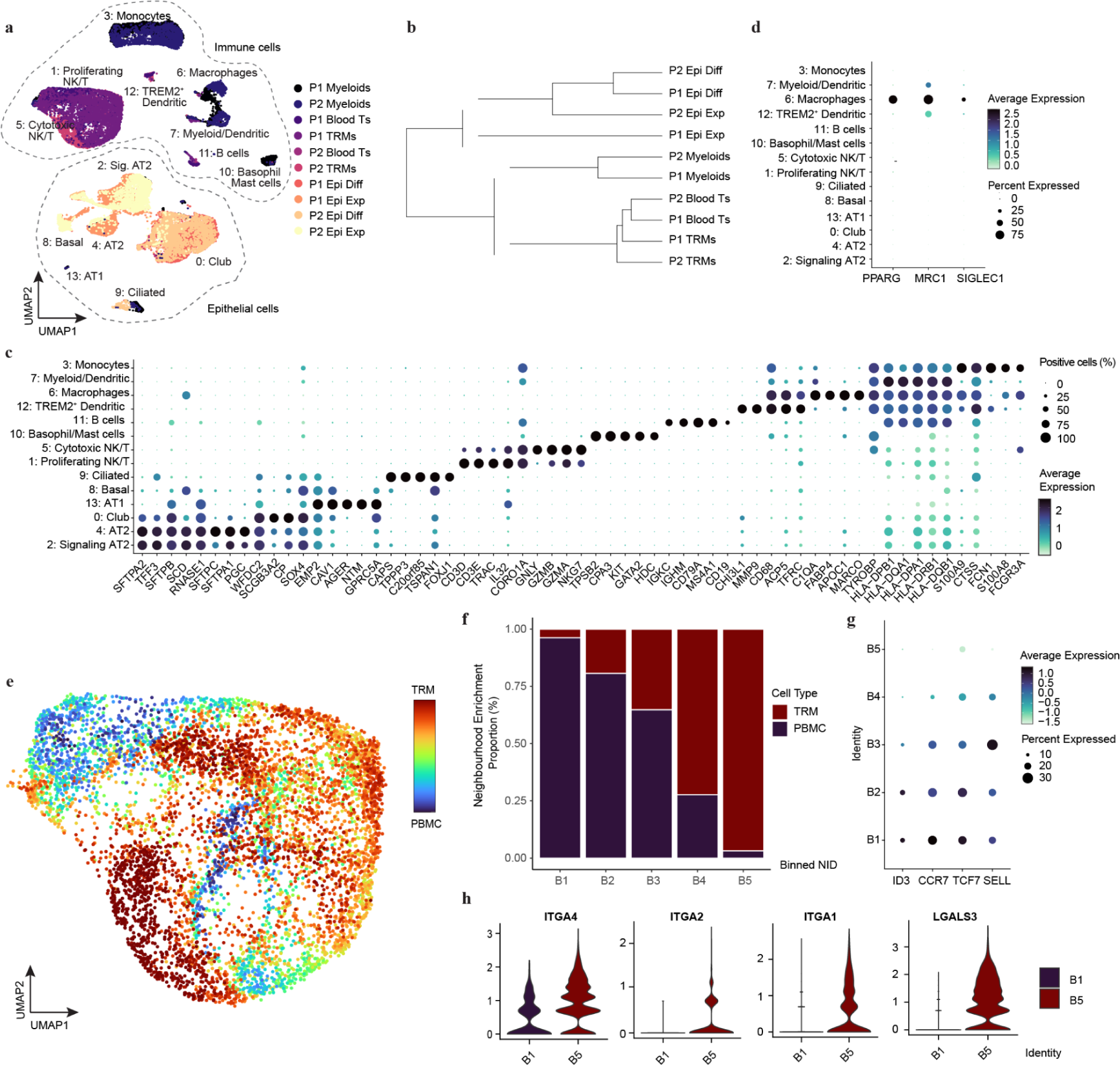
Consistent donor profiles and distinct differences between TRMs and blood-derived T cells. **a**, Integrated UMAP embedding from scRNA-seq data of patient-derived lung cells coloured and labeled by sample. **b**, Clustering dendrogram of samples based on their Euclidean distance. **c**, Dotplot summarizing the expression patterns of representative genes (x-axis) across the clusters (y-axis). **d**, Expression of *PPARG*, *MRC1* and *SIGLEC1* across all clusters. Dotsize reflects positive population (%) while color represents average expression. **e**, UMAP embedding of TRMs and PBMCs. Colour scale reflects neighbourhood enrichment of TRMs or PBMCs in individual cells. **f**, Stacked barplot showing proportions of TRMs and PBMCs in 5 equidistant bins of the neighbourhood enrichment score. **g**, Expression of PBMC representative genes *ID3*, *CCR7*, *TCF7* and *SELL* across the different bins introduced in **f**. **h**, Violin plots of TRM marker genes (*ITGA4*, *ITGA2*, *ITGA1*, *LGALS3*) shown for bin 1 (B1) and bin 5 (B5).

**Extended Data Figure 3:**
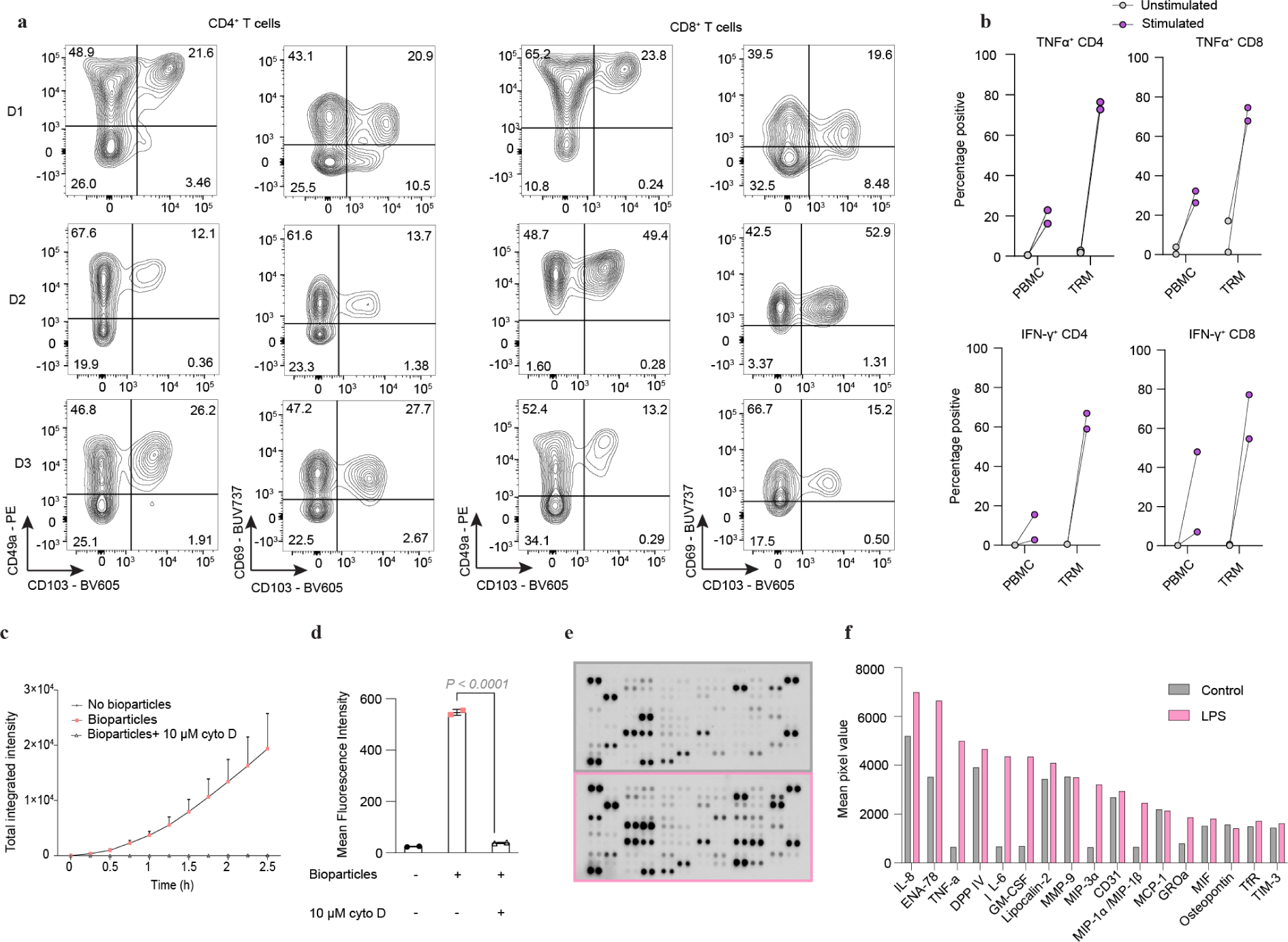
Characterisation and functional analysis of TRMs and alveolar myeloid cells after cryopreservation. **a**, Flow cytometry plots of T cells separated into CD4+ (left) and CD8+ (right) populations. Gated on viable, single, CD45+ CD3+ cells. The different rows represent the results for different donors (D). The left column shows the CD49a - PE signal versus CD103 - BV605, the right column shows the CD69 - BUV737 signal versus CD103 - BV605. **b**, Line graph showing the TNFα-positive (top row) and IFNy-positive (bottom row) percentages in CD4+ and CD8+ T cells at baseline or upon stimulation for PBMC and TRM. The dot represents the mean of technical replicates for each biological replicate. Data shown for two independent biological replicates. **c**, Line graph showing the total integrated intensity over the course of culturing the alveolar myeloid cells with or without *S. aureus* bioparticles for over 2.5 h. Cytochalasin D (cyto D) was added as a phagocytosis inhibitor. Technical duplicates, data shown of one representative biological donor. Mean and SD. **d**, Barplot presenting the mean fluorescence intensity after 3 h of culturing the alveolar myeloid cells with and without the bioparticles. One-way ANOVA with Tukey’s multiple comparisons test. **e**, Images of proteome profiler blots shown for the control in grey and for LPS treatment in pink. The different dots indicate different cytokines measured from supernatants of overnight treated alveolar myeloid cells. **f**, Barplots representing the mean pixel value for the two blots shown in **e**. The highest expressed proteins are presented in the plot. Data of one independent biological replicate shown.

**Extended Data Figure 4:**
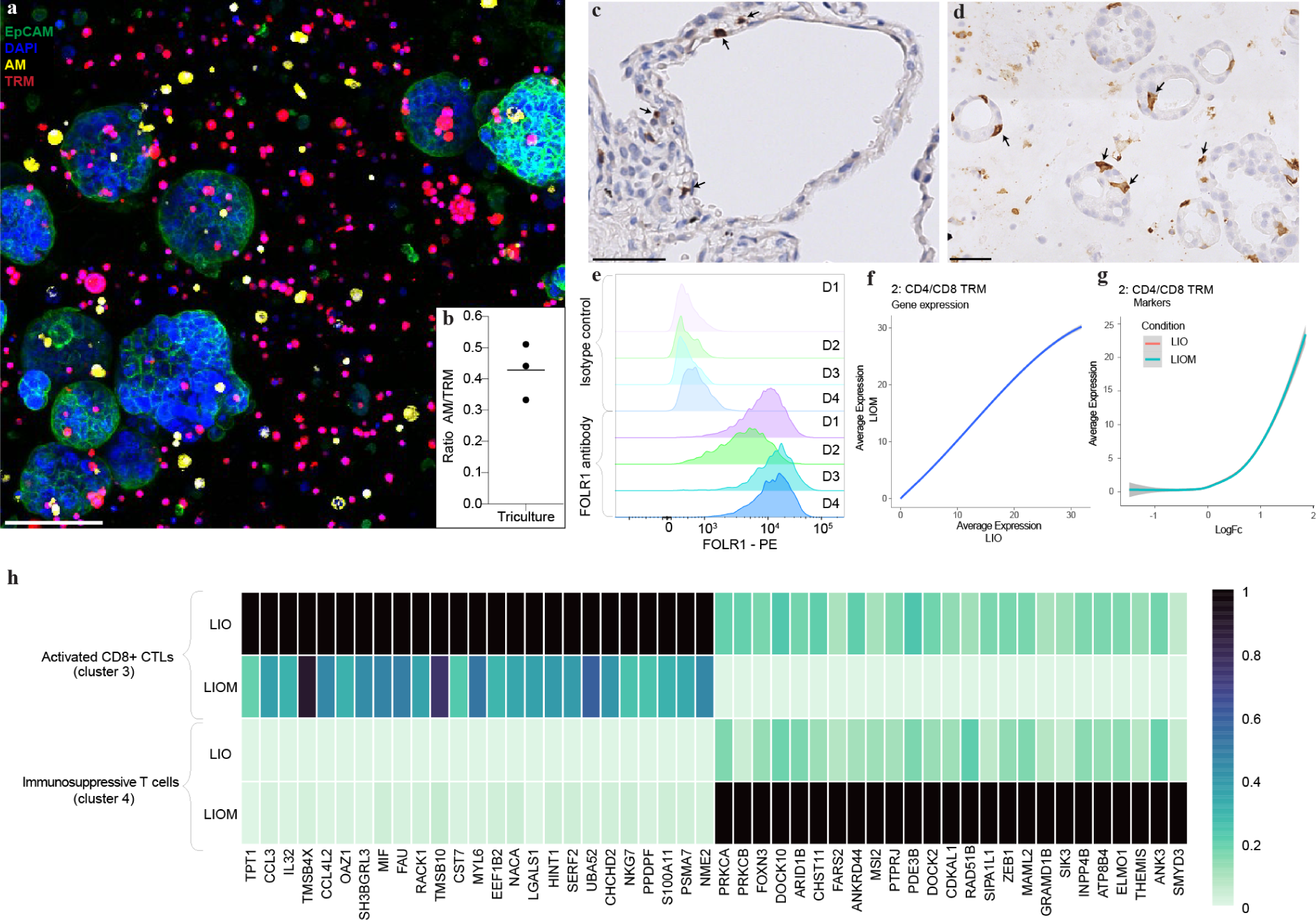
Comprehensive characterisation of the in vitro model and the effect of alveolar myeloid cells on TRMs. **a**, Immunofluorescent image of a live LIOM culture with labelled TRMs (red) and AMs (yellow), nuclei (blue) and epithelial cells (EpCAM - green). Scale bar, 100 µm. **b**, Quantifications of **a**, showing the ratio of AMs to TRMs, line at mean. Dots represent technical triplicates. **c**, IHC staining of CD103 (brown) on FFPE lung tissue. Arrows indicate CD103+ cells. Scale bars, 50 µm. **d**, IHC staining of CD103 (brown) on FFPE LIOM samples. Arrows indicate CD103+ cells. Scale bars, 50 µm. **e**, Histograms of flow cytometry analyses of alveolar organoids showing the isotype control and the FOLR1 staining for four independent biological replicates. Gated on viable, single EpCAM+ cells. **f**, Transcriptome-wise average gene expression of TRMs in LIO (x-axis) and LIOM (y-axis) cultures. **g**, Log-normalized fold change (x-axis) and average expression (y-axis) of TRM markers in LIO and LIOM cultures. **h**, Heatmap of marker gene expression for cluster 3 (cytotoxic CD8+ T cells) and 4 (immunosuppressive T cells) for LIO and LIOM cultures.

**Extended Data Figure 5:**
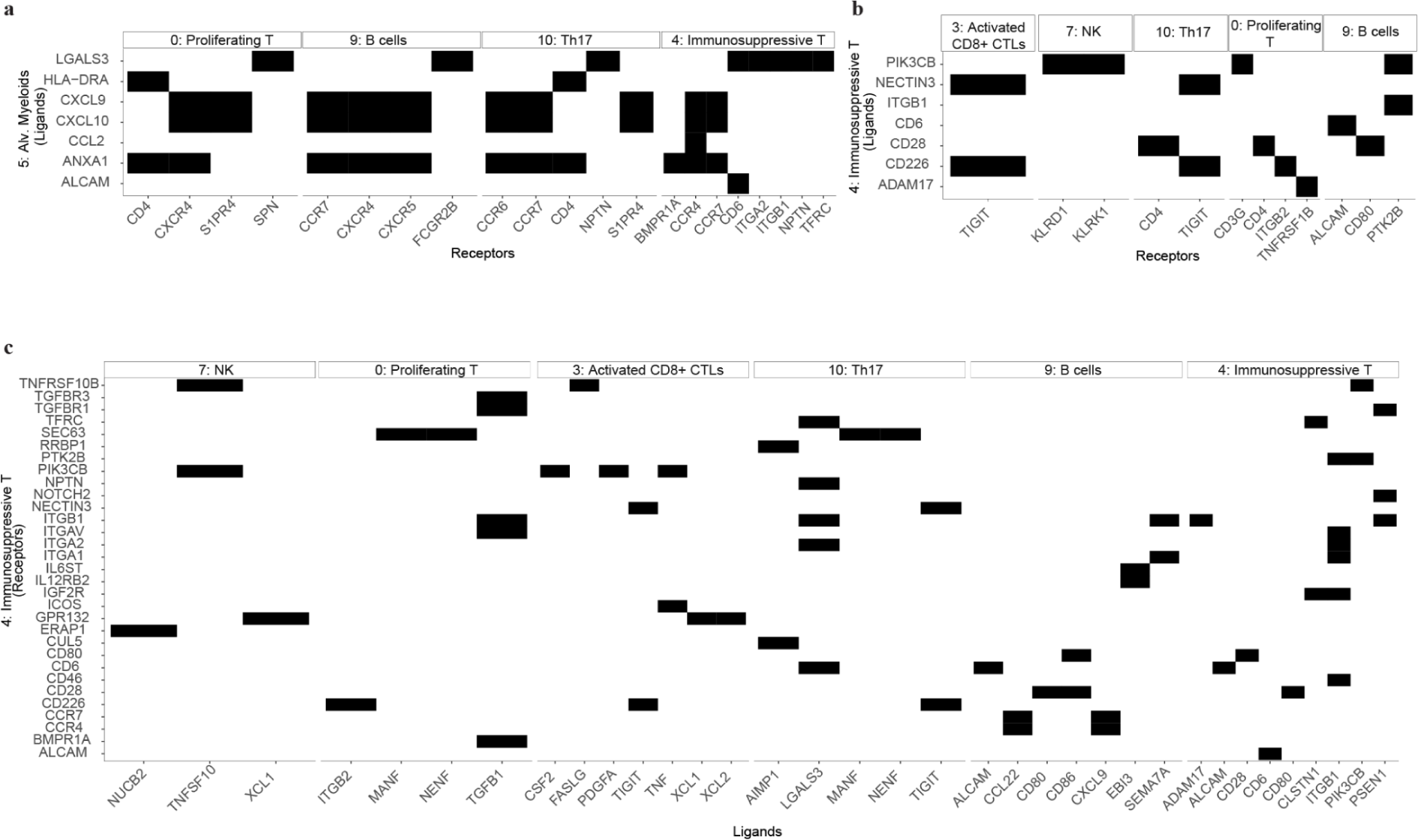
Prediction of interactions between secreted ligands and receptors across different cell types. **a**, Interaction map between alveolar myeloid secreted ligands (rows) and receptors (columns) expressed on other cell types. Plot is coloured based on binary presence-absence of detected interaction. **b**, Interaction map between immunosuppressive T cell secreted ligands (rows) and receptors (columns) expressed on other cell types. **c**, Interaction map between immunosuppressive T cell receptors (rows) and ligands (columns) expressed on other cell types.

**Extended Data Figure 6:**
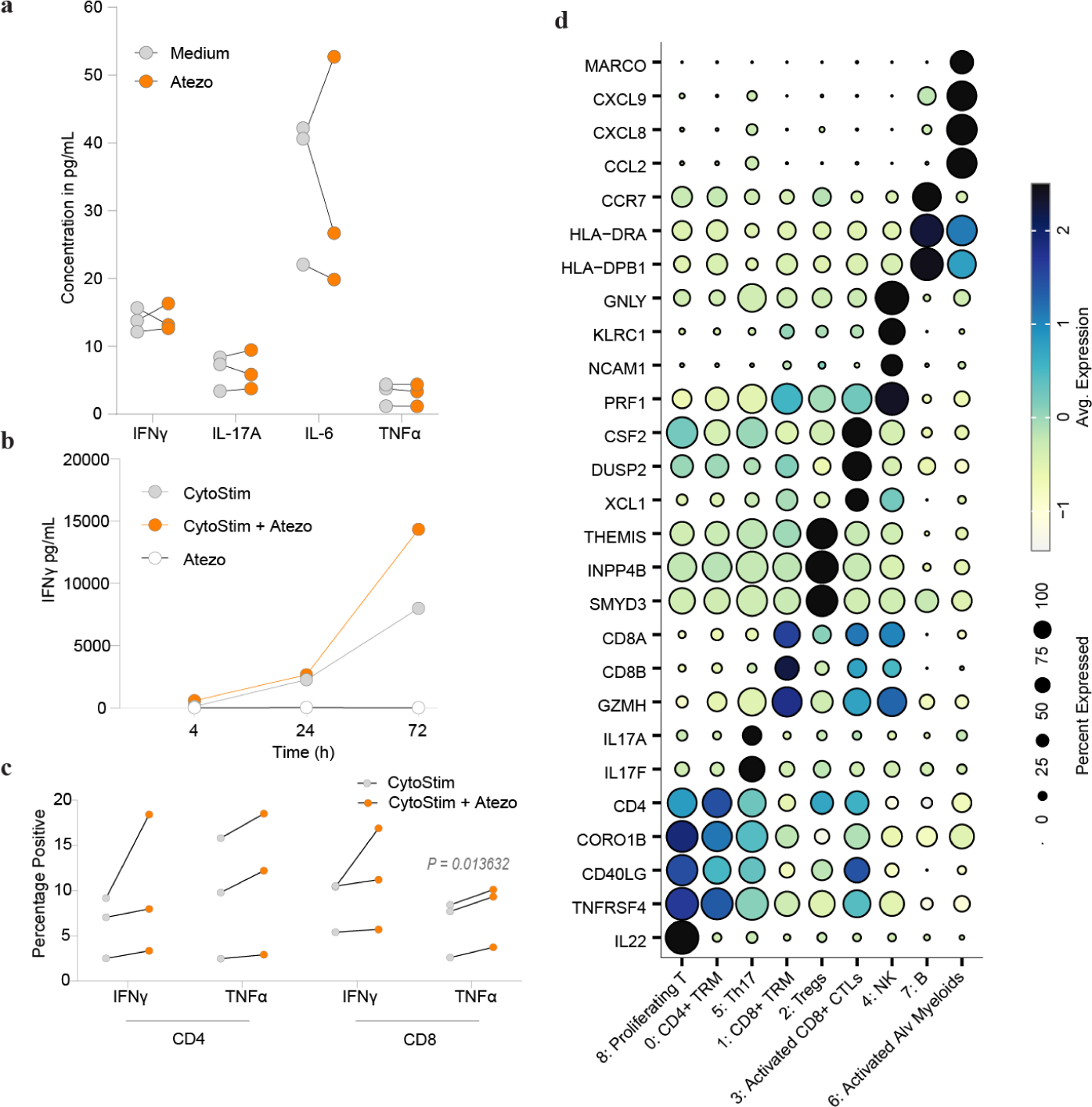
Atezolizumab-treated LIOMs increase intracellular and secreted cytokines upon T cell stimulation. **a**, Lineplots showing the concentration of four cytokines (IFNγ, IL-17A, IL-6, TNFα) measured in the supernatants of untreated and atezolizumab-treated LIOMs after 72 h of culture. Three biological independent replicates. Atezolizumab at 10 µg/mL. **b**, Lineplot of IFNγ concentrations at 3 different time points (4, 24, 72 h) measured in the supernatant of atezolizumab-treated LIOMs in the presence or absence of CytoStim. One representative biological donor. **c**, Lineplots showing the differences of intracellularly measured cytokines (IFNγ and TNFα) by flow cytometry in CytoStim or CytoStim + atezolizumab-treated LIOMs after 72 h. **d**, Dotplot summarising the most highly enriched genes across clusters as identified in Fig. 4f.

## Methods

### Human samples

Normal tissue adjacent to tumour human lung samples and annotated data were obtained upon informed patient consent, and all experiments were performed as defined in the framework with BeCytes (Barcelona, Spain). The lung samples were collected as part of the project “Modelling cancer-immunotherapy induced pulmonary toxicities with in vitro human assays” Code P/22-002/MCIPT.2022.v1”, approved by the Research Ethics Committee on Medicinal Products (CEIm) of the Fundació Assistencial Mútua Terrassa in ordinary session #02/2022 on February 23, 2022, and by the Research Ethics Committee on Medicinal Products (CEIm) of Parc Taulí de Sabadell (Barcelona) in ordinary session #2022/5030 on March 29, 2022. 20 consenting patients underwent tumor resections with partial resection of lung tissue and for our experiments we used micro-and/or macroscopically tumour-free regions.

### Lung tissue preparation

Lung tissue was first processed, by washing it thoroughly several times with the 1 x DPBS buffer containing penicillin [1500U/ml] (Thermo Fisher), streptomycin [1500 pg/ml] (Thermo Fisher), gentamicin [500 pg/ml] (Thermo Fisher) and amphotericin [12.5 µg/ml] (Thermo Fisher). The pleura was removed and the tissue cut into smaller pieces for the subsequent isolation of the different cell types.

### Isolation of TRM cells

A part of the washed lung tissue was used to isolate TRM with the scaffold-based egression method (adapted from ref.^25,26^). Small explants were placed onto a 9 x 9 x 1.5 mm^3^ tantalum-coated carbon-based scaffold (Ultramet). The scaffolds were then transferred to a well of a collagen I - coated 24-well plate (Corning) with 1 mL/well of cytokine-containing immune cell medium (10 IU/mL IL-2 (Roche), 2 ng/mL recombinant human IL-15 (BioLegend), RPMI 1640, 10% FBS, 1500 U/mL penicillin, 1500 pg/mL streptomycin, 500 pg/mL gentamicin, 12.5 pg/mL amphotericin B) and were cultured for 7 days. After 3 days of culture, 1 mL of medium was added to the cells. After 4 more days, the scaffolds with the tissue were removed from the well and the cells in the well were harvested via pipetting.

### Alveolar myeloid cell isolation

A piece of the washed lung tissue was used to isolate the alveolar myeloid fraction (adapted from ref. ^55^). In brief, the lung tissue was punctuated with a syringe (0.8×40 mm) and 0.1 M NaCl (Thermo Fisher) was injected several times to flush out the cells. Once the tissue was free of blood, the cell suspension was harvested, centrifuged and the alveolar myeloid cells were negatively selected with the EasySep human monocyte enrichment kit without CD16 depletion (STEMCELL Technologies) according to the manufacturer’s instructions.

### Alveolar type 2 isolation for alveolar organoid generation

Roughly 5 g of washed tissue was used to isolate alveolar epithelial type 2 (AT2) cells as described previously by Katsura et al^10^. In short, the cleaned tissue was minced into very small pieces, using scissors and a scalpel, and transferred to a C tube (Miltenyi Biotec) containing RPMI 1640 with 1.68 mg/mL collagenase I (Thermo Fisher), 10 U/mL DNAse (Roche) and 5 U/mL Dispase (Corning). The C tube containing the minced tissue was inserted onto the gentleMacs™ Octo Dissociator (Miltenyi Biotec) with heater and the 37C_h_TDK_1 programme was started. Upon programme completion, the cell suspension was filtered through a 70 µm filter (Miltenyi Biotec) and centrifuged. The cell pellet was resuspended in anti-human mouse IgM HT2-280 antibody (Terrace Biotech) and Human TruStain FcX (BioLegend) for 1 h at 4 °C, followed by the addition of anti-mouse IgM microbeads (Miltenyi Biotec) for another 15 min at 4 °C. The cells were positively selected with LS columns (Miltenyi Biotec) on a OctoMACS™ Separator (Miltenyi Biotec). HT2-280^+^ cells were counted manually using a C-chip and resuspended at 1 million cells/mL in Matrigel Matrix GFR (Corning) and 20 µL domes were seeded onto a tissue-culture treated 24-well plate (Corning). The plate was flipped and incubated for 15 min at 37 °C. After incubation, PneumaCult™ Alveolar Organoid Expansion (AvOE) Medium with 1x of seeding supplement (STEMCELL Technologies) and 1:250 primocin (InvivoGen) was added to the domes.

### Alveolar organoid culture

The alveolar organoids were kept in PneumaCult™ AvOE Medium Medium (STEMCELL Technologies) with media changes 2 times per week for roughly 2 weeks before they were split. For splitting, Gentle Cell Dissociation Reagent (STEMCELL Technologies) was added to the domes for 10 min and then the cell suspension was centrifuged. The organoid pellet was disturbed by rough pipetting with DMEM F-12 (ThermoFisher) + 1% BSA buffer and another centrifugation step. The organoid fragments were resuspended in Matrigel Matrix GFR (Corning) at a split ratio of 1:3 and seeded in 20 µL droplets onto 24-well plates. The plate was flipped and incubated for 15 min before PneumaCult™ AvOE Medium containing 1x of passage supplement was added to the domes and left for 3-4 days. In order to differentiate the alveolar organoids, they were grown for 10 days in PneumaCult™ AvOE before the medium was replaced with PneumaCult™ Alveolar Organoid Differentiation Medium for another 10 days with 2 media changes per week.

### Preparation of LIOs and LIOM, including treatments

Alveolar organoids were grown in expansion medium and used at passage 3. On the day of the co-culture setup, the alveolar organoids were harvested with ice cold Cell Recovery Solution (Corning) for 40 min. Organoids were collected and centrifuged. Donor-matched immune cells (TRMs and AMs) were thawed and counted. For LIO cultures, organoids were combined with TRMs. A 20 µl organoid dome was harvested and resuspended in 15 µl of matrix, whereas TRMs were used at a density of 10,000 cells mm^−3^ of resuspension volume. For LIOM cultures, in addition to the LIO, alveolar myeloid cells were added at 4,000 cells mm^−3^. After combining the cells, the suspension was centrifuged and resuspended either in Matrigel Matrix GFR (Corning) or, if cultures were supposed to be fixed with formalin, in a 1:1 mixture of Matrigel Matrix GFR and 4 mg ml−1 Cultrex Rat Collagen I (R&D Systems). For time-lapse imaging and live imaging experiments, the TRMs were labelled with CellTrace^TM^ Far Red (ThermoFisher) and the alveolar myeloid cells with CFSE (ThermoFisher) before the immune cells were combined with the organoids. After plating the LIOs and LIOMs, a 50:50 mix of RPMI 1640 10% FCS and PneumaCult™ AvOE Medium was used for cultures.

To assess T-cell induced inflammation, the FOLR1 TCB (Roche or Creative Biolabs) and its non-targeting control (one arm contains a CD3 binder and a non-specific DP47 arm) were added directly to the culture medium after co-culture setup. The concentration used was 1 ng/mL. To treat the cultures with known pneumonitis-inducing drugs, erlotinib (Selleckchem) was used at 1 µM and atezolizumab (Roche) was used at 10 µg/mL. To stimulate the T cells, CytoStim (Miltenyi Biotec) was added to the medium at 20 µL/mL. The immune cells were stained with CellTrace^TM^ Far Red and CellTrace^TM^ Yellow Cell Proliferation Kit (ThermoFisher) according to the manufacturer’s instructions by using them at a final working concentration of 1 µM.

### Cytokine production analyses of T cells by flow cytometry

T cells (either from tissue or the blood) were seeded at 100,000 cells/well of an ultra-low attachment 96-well plate (Corning). The medium was either supplemented with 1x of Cell Stimulation Cocktail (ThermoFisher) or 1x Protein Transport Inhibitor Cocktail (ThermoFisher). Following treatment, the cells were incubated for 4 h at 37 °C and then stained for surface proteins (**Supplementary Table**). The cells were fixed and permeabilized using the FoxP3 Transcription Factor Staining Buffer Set (ThermoFisher). Subsequently, intracellular proteins were stained before acquisition on a BD Fortessa X-20 using BD FACSDiva Software v.9.7 and analysed using FlowJo v.10.

### Lactate dehydrogenase (LDH) measurements

To assess treatment-induced toxicity, the supernatants were directly used after sampling to measure the lactate dehydrogenase (LDH) content. The cytotoxicity kit (Roche) was utilized according to the manufacturer’s manual. In short, a standard curve was prepared and the samples were diluted in PBS, combined with the reaction mix and incubated on a shaker in the dark for 30 min. The plate was read on a PerkinElmer Envision, 2104 Multilabel reader with absorbance at 490 nm.

### Epithelial cell cytotoxicity assessment with Caspase 3/7

LIO and LIOM cultures were prepared as described above and seeded in 5 µL matrigel domes onto a PhenoPlate™ 96-well microplate (Revvity). Apoptosis was assessed by measuring the CellEvent Caspase 3/7 Detection Reagent (Invitrogen) intensity. The CellEvent Caspase 3/7 reagent was added to the medium at 1:1,000. The wells were imaged as often as every 2 hours using the Operetta CLS (Revvity) with a 5x air objective in confocal mode. Images were acquired using a z-stack of roughly 450 µm, depending on the cultures. The distance between each stack was set to the minimum of 26 µm for the employed objective (autofocus: two Peak; binning: 2). Brightfield and Alexa Fluor™ 488 were selected as channels and each well was imaged entirely by selecting 4 fields. The plates were kept inside of the machine for the duration of the time course and the temperature was set to 37 °C with 5% CO_2_. Caspase 3/7 fluorescence signal intensity was analyzed with the Opera Harmony software v.4.9 (Revvity). In the analysis workflow, the organoids were first segmented using the ‘Find Texture Regions’ based on the brightfield signal, then selected and identified as single organoids with the options ‘Select Region’ and ‘Find Image Region’. Following the identification of the objects, the morphology parameters of interest (>10000 µm^2^) were defined with ‘Calculate Morphology Parameters’ followed by ‘Select Population’. Lastly, the Caspase 3/7 signal was quantified per individual organoid with the ‘Calculate Intensity Properties’ of the Alexa Fluor™ 488 channel.

### Cytokine analysis of LIO and LIOM cultures

The supernatants of the cultures were collected and directly frozen down and stored at −80 °C. For the measurements, two different multiplex custom-made ProcartaPlex immunoassay kits were purchased from invitrogen (PPX-10-MXFVMZC (GM-CSF, GzmB, IFN-γ, IL-2, IL-4, IL-6, IL-10, IP-10, MCP-1, TNF-α) and PPX-7-MXPRM6K (IL-2R, IL-1RA, IL-6, IL-10, IL-17 TNF-α, IFN-γ)) and used according to the manual. In short, the supernatants were diluted 1:10 with the universal assay buffer, the standards prepared, the magnetic capture beads added to the plate and washed with the automatic plate washer (405TS microplate washer from Bioteck). Samples and standards were then added to the microbeads and incubated on a shaker for 2 h at RT in the dark. Following the incubation, the beads were washed and the detection antibodies were added to the beads for 30 min at RT in the dark. Again, the beads were washed and Streptatividin was added for another 30 min at RT in the dark. After the incubation time, the microbeads were washed and they were resuspended in the reading buffer. The samples were measured with the Bioplex-200 instrument from Bio-Rad with the corresponding Bio-Plex Manager Software version 6.2.

### Fixation and embedding of in situ collagen domes

In situ embedding was done according to previously published protocols^25,56^. In short, cultures were plated in a 1:1 Matrigel Matrix GFR (Corning) and 4 mg/mL Cultrex Rat Collagen I (R&D Systems) mix. At the end of the culture, the domes were washed with 1x DPBS followed by a fixation step with 4% paraformaldehyde (PFA) solution (Thermo Fisher) for 30 min at RT. After fixation, the domes were washed twice with 1x DPBS and kept in 1x DPBS at 4 °C until used for the next step. Epredia™ HistoGel™ Specimen Processing Gel (Thermo Fisher) was warmed up in the microwave and kept at 64°C until usage. 400 µL of histogel was added to the domes, left to polymerize for 20 min at 4 °C before gently removing the Histogel from the well using a bent spatula. The embedded samples were then transferred to biopsy cassettes and consequently dehydrated in the HistoCore PEARL tissue processor (Leica) overnight. Following dehydration, the samples were embedded in liquid paraffin. Generated FFPE blocks were cut on a Microtome (Thermo Fisher) with a thickness of 5 µm and the slices were transferred to Epredia™ SuperFrost Ultra Plus™ GOLD Adhesion Slides (Thermo Fisher). The slides were baked at 50 °C for up to 48 h or until dry.

### Hematoxylin and eosin (H&E) stainings

Slides were deparaffinized and hydrated to distilled water before staining with hematoxylin (Sigma-Aldrich) for 5 min followed by a rinsing step under running water for 5 min. Then, the slides were washed with 0.3% acid alcohol for 20 s and another additional 5 min wash under running water. The slides were subsequently counterstained with eosin (Thermo Fisher) for 2 min, dehydrated, cleared and mounted.

### Immunohistochemistry (IHC) stainings with the Ventana Discovery Ultra

The dried slides containing the slices from the FFPE were stained using a Ventana Discovery Ultra (Roche Tissue Diagnostics). See **Supplementary Table** for used antibodies and dilutions. Firstly, the slides were baked at 60°C for 8 min followed by 3 cycles of deparaffinization with the Discovery Wash at 69°C, 8 min for each cycle. The slides were then pretreated with Discovery CC1 for 40 min at 92°C followed by an incubation with the Inhibitor CM (ChromoMap DAB kit, Roche) for 4 min, RT. Subsequently, primary antibody was added at the indicated dilutions and incubated for 60 min at 37°C. Following the primary antibody incubation, the appropriate secondary antibody (multimer HRP) was added for another 16 min, RT. The secondary antibody was then labeled with DAB by incubating the slides for 8 min. Lastly, the nuclei were counterstained with hematoxylin for 8 min and post counterstained with bluing reagent for 4 min. The slides were cleared in xylene and mounted.

### Multiplex immunofluorescence (mIF) staining and scanning

The mIF staining was performed as described previously^56^. In short, the Ventana Discovery Ultra Automated Tissue Stainer (Roche Tissue Diagnostics) was applied for the mIF staining of FFPE cuts. See **Supplementary Table** for used antibodies and dilutions. Slides were baked for 8 min at 60 °C followed by a heating step at 69 °C for 8 min and deparaffinized with the Discovery Wash. This cycle was repeated twice and afterwards antigens were retrieved at 92 °C with Tris-EDTA buffer (pH 7.8, Ventana) for 40 min. A Discovery Inhibitor (Ventana) was applied for 16 min for blocking and then neutralized. Primary antibodies were diluted in 1x Plus Automation Amplification Diluent (Akoya Biosciences) and the slides were incubated with these antibodies for 40-60 min. Primary antibodies were detected using anti-species secondary antibodies conjugated to HRP (OmniMap Ventana). The secondary antibody incubation time was 12 min followed by another 12 min incubation with the relevant Opal dye (Akoya Bioscience). After every cycle of incubation with primary and secondary antibody and opal dye, an antibody neutralization and HRP-denaturation step were applied to remove residual antibodies as well as HRP, before starting a new cycle with the Discovery Inhibitor blocking step. In the last sequence, Opal TSA reagent was applied instead of the Opal dye followed directly by the addition of Opal dye 780 for 1 h incubation. Lastly, samples were counterstained with 4’,6-Diamidino-2-phenylindol (DAPI, Roche). mIF stainings were scanned with multispectral imaging by the Vectra® Polaris™ (Akoya Biosciences) using the MOTiF™ technology at 20x magnification for all colors (Opal 480, Opal 520, Opal 570, Opal 620, Opal 690, Opal 780 and DAPI). To reduce variation between slides, they were acquired as a batch ensuring the same setting and therefore the comparability for subsequent image analysis. PhenoChart (v1.0.12) and inForm (v2.4) were used for the unmixing of the channels and tiling of the images. Tiles were fused in HALO AI (Indica labs, v3.2.1851.328).

### Image analysis with HALO AI of IHC

HALO AI (Indica Labs, v3.2.1851.328) was used to analyze single IHC stained images. For the single stainings, three distinct regions were defined by a board certified pathologist per donor: healthy, inflamed and inflamed with black pigment in macrophage aggregates. The same regions of each donor were used on the subsequent serial section to analyze the staining. For the analysis, the Multiplex IHC version 3.4 module was used to perform nuclear segmentation based on DAPI+ cells. Only the specific staining for the different cell types were analysed on DAPI+ cells. Background was added as an exclusion marker to optimize the nuclear and specific marker detection. Percentages of positive cells were obtained by dividing the number of positive cells by the total of cells cleaned. Furthermore, the cells were quantified per area.

### Time-lapse imaging of LIO culture

LIO were prepared as described above with TRMs stained with CellTrace^TM^ Far Red. Time lapse-imaging was performed 24 h post co-culture set-up with Leica a STELLARIS 8 confocal microscope using a dry objective (HC PL APO ×10/0.40 CS2) and 0.75 zoom. Images were obtained in bidirectional mode with 1,024 × 1,024 pixels at 600 Hz. Images were acquired with a 2.5 µm z-stacks. Samples were imaged for 60 min while being kept in an incubation chamber (The Box, Life Imaging Services) at 37 °C and 5% CO2. Following acquisition, maximum-intensity projections were generated with Leica Las X software and later exported as AVI files using ImageJ v.1.54i.

### 3D immunophenotyping of LIOMs

Immune-organoid co-cultures were set up as described above. The cell pellet was resuspended in a 1:1 mix of Matrigel Matrix GFR and TeloCol^®^-6 bovine solution, 6 mg/mL (Advanced BioMatrix) and 20 µL domes were seeded onto 6-well tissue-culture treated plates (Corning). After culturing the cells, the domes were washed with 1x DPBS and fixed after 1x BD Cytofix (BD BioSciences) at 4 °C for 20 min. The domes were subsequently washed 3 times with 1x DPBS and kept in 1x DPBS at 4°C until the next step where the domes were embedded in 4% low gelling temperature agarose (Sigma) and sectioned into 100 µm sections with a Vibratome (Leica 1200s). Sections were subsequently stained with panels of fluorescently conjugated antibodies (see **Supplementary Table**), cover-slipped with Fluoromount G mounting media (SouthernBiotech), and imaged on a Leica SP8 microscope using 40×1.3 NA (HC PL APO 40x/1.3 Oil CS2) oil objective with type F immersion liquid (Leica).

### Imaris image analysis of 3D immunophenotyping images

Imaris (Oxford Systems, version 10.1.1) was used for image visualization and analysis. Downstream image analysis, such as phenotyping and cell type enumeration, was performed in Matlab 2022a. Channel arithmetic was performed using the default Imaris function in order to remove channel bleed-through. Imaris was used to segment individual cell objects that represent different cells. After surface creation, the MFI for each imaged channel, as well as the volume, sphericity, and position of the cell objects were exported and concatenated into a single.csv file using the Imaris_To_FlowJo_CSV_Converter_V6 MATLAB function, available online. The combined.csv file was next imported into FlowJo (TreeStar) and the cell objects were classified into the indicated cell subsets according to the gating strategies shown in the respective figures.

### Cytospin of alveolar myeloid cells

Cryopreserved alveolar myeloid cells were thawed, centrifuged and the cell pellet was resuspended at 500,000 cells/mL. Superfrost Plus adhesion microscope slides (Epredia) were inserted into the holder. The filter card (Thermo Fisher) was put on the slide and the cytofunnel (ThermoFisher) positioned next to it before the holder was fastened. 70 µL of the cell suspension was loaded into the cytofunnels and the slides were centrifuged in the Cytospin 4 Centrifuge (Thermo Fisher) at 1000 rpm for 5 min. Then the cells on the slides were fixed with 4% PFA and stored at −80°C before further H&E staining.

### Assessments of phagocytic activity with *S. aureus* bioparticles

Cryopreserved alveolar myeloid cells were thawed, counted and plated at 30,000 cells/well of a PhenoPlate™ 96-well microplate (Revvity) in RPMI 1640 with 10% FBS, 1 mM sodium pyruvate (invitrogen), 1x HEPES (Thermo Fisher), 50 ng/mL M-CSF (Miltenyi Biotec), 500 U/mL penicillin, 1500 pg/mL streptomycin (Thermo Fisher), 500 pg/mL gentamicin (Thermo Fisher), 12.5 pg/mL amphotericin (Thermo Fisher). The cells were incubated at 37 °C overnight. The following day, pHrodo™ Red *S. aureus* BioParticles™ Conjugates for Phagocytosis (Thermo Fisher) were prepared according to the manual and added at 1:10 to the culture medium. The conditions that received cytochalasin D to inhibit the phagocytic capacity were incubated with 10 µM of cytochalasin D (Sigma-Aldrich) for 1 h at 37°C prior to the addition of the beads. The plate was imaged with the standard programme using a 20x air objective in the Incucyte® S3 Live-Cell Analysis Platform (Sartorius) every 20 min for 3 h. After the final image acquisition, the plate was imaged with the Operetta (Revvity) with a 20x Water Objective and the data was analysed using the Harmony software (Revvity).

### Proteome profiler analysis of alveolar myeloids

Alveolar myeloid cells were plates at 500,000 cells per well of a collagen-I coated 6-well plate (Corning) in RPMI 1640 with 10% FBS, 1 mM sodium pyruvate (Invitrogen), 1x HEPES (Invitrogen), 500 U/mL penicillin, 1500 pg/mL streptomycin, 500 pg/mL gentamicin (Thermo Fisher), 12.5 pg/mL amphotericin (Thermo Fisher). 100 ng/mL of LPS (Thermo Fisher) were applied to the cells for 24 h to induce the cytokine production. After 24 h, the supernatants were collected and stored at −80°C. Supernatants were then analyzed with the Proteome Profiler human XL cytokine array kit (ARY022B, R&D systems). Proteome analysis was done according to the manufacturer’s instructions. Briefly, cell supernatants were diluted and incubated with the array membranes overnight at 4 °C on a rocking platform shaker. The next day, arrays were washed in order to remove unbound proteins followed by the incubation with the detection Antibody Cocktail for 1 h at room temperature on a shaker. The membranes were washed again and 1x Streptavidin-HRP was added for another 30 minutes while shaking. A last wash was done before the chemiluminescent detection reagent was added to the membranes.The signals were detected by a ChemiDoc MP system and quantified by ImageLab v. 6.0.1.

### MHC-II-Associated Peptide Proteomics (MAPPs) Assay

Stock solutions of the highly immunogenic antigen keyhole limpet hemocyanin (KLH-Imject Maleimide-Activated mcKLH; Thermo Fisher), which served as a positive control, were reconstituted in sterile water at 10 mg/mL and stored at 4 °C. MAPPs assay was performed according to the recently published protocol^57^ and analyzed according to Steiner et al^58^. In short, myeloid cells were challenged with the positive control KLH (50 μg/mL), and matured with lipopolysaccharide (LPS; 1 μg/mL) from *Salmonella enterica* (Sigma Aldrich) for 24 h at 37 °C and 5% CO_2_. Mature cells were harvested, washed with PBS and lysed in a hypotonic buffer (20 mM Tris-HCl pH 7.8; 5 mM MgCl_2_) containing 1% (v/v) Triton X-100 Surfact-Amps detergent solution (Thermo Fisher) and a protease inhibitor mini tablet (Thermo Fisher) for 1 h in a ThermoMixer (Eppendorf) at 1100 rpm and 4 °C. Lysates were then collected and frozen at −80 °C until immunoprecipitation. Lysates were incubated with the antibody clone L243 (10 μg; anti-human HLA-DR biotin; RayBiotech) overnight at 4 °C on a rotator. Immunoprecipitation of MHC-II receptors was carried out using the automated AssayMAP Bravo platform (Agilent Technologies) and accompanying streptavidin cartridges (Agilent Technologies). HLA II-specific peptides were eluted in 0.1% (v/v) TFA in a final volume of 18 μL. Peptide samples were then directly loaded onto Evosep C18 tips (Evosep Biosystems) according to the manufacturer’s recommendations and stored at 4 °C until LC-MS/MS analysis (details can be found in ref ^57^). Detected peptides were grouped into clusters and represented along the KLH sequence using dataMAPPs, an in-house-developed R-based workflow^58^.

### Immunofluorescence stainings of alveolar organoids and alveolar myeloids

Alveolar organoids were cultured and differentiated as described above. The organoids were harvested using Cell recovery solution (Corning) and fixed with 4% paraformaldehyde for 30 min. Alveolar myeloids were thawed and seeded onto a PhenoPlate™ 96-well microplate (Revvity) and left to attach overnight at 37°C before being fixed with 4% PFA for 30 min. After fixation, the samples were washed three times with PBS before a blocking step with blocking buffer (5% BSA, 0.3% Triton X-100 in 1 x DPBS (+/+)), followed by the addition of the primary antibodies overnight (see **Supplementary Table**). The next morning, samples were washed with 1% BSA, 0.3% Triton X-100 in 1x DPBS (+/+) and then incubated with secondary antibodies and Nucblue (Thermo Fisher) overnight. After the incubation, the cells were again washed with 1% BSA, 0.3% Triton X-100 in 1x DPBS (+/+) and organoids were transferred to a PhenoPlate™ 96-well microplate (Revvity) in 1x DPBS. The samples were imaged with a 20x air objective (HC PL APO 20x/0.75 CS2) and a 63x oil-immersion objective (HC PL APO 63x/1.40-0.60 OIL) on the Leica STELLARIS 8 confocal microscope. The images were acquired using the bidirectional mode with 1024 x 1024 pixels at 600 Hz with z-stacks of different ranges using the optimal z-stack size. Displaying images was done using the Leica Las X software and ImageJ.

### Live imaging of LIOM

LIOM cultures were set up as described above. Immune cells were labelled either with CellTraceFar Red Cell Proliferation Kit (Thermo Fisher) or the CellTrace^TM^ Yellow Cell Proliferation Kit (Thermo Fisher) according to the manufacturer’s instructions. LIOMs were seeded in 5 µL domes onto a PhenoPlate™ 96-well microplate (Revvity). After the domes solidified at 37°C for 15 min, the epithelial cells were stained with the primary-labeled CD326 (EpCAM) Monoclonal Antibody (323/A3), Alexa Fluor™ 488 (Thermo Fisher) in 1x DPBS at 1:50. Additionally, Hoechst 33342 Solution (BD BioSciences) was added to the staining mix at 1:2,000 and the cells were incubated for 30 min at 37°C. Following the incubation, the domes were washed with 1x DPBS and directly imaged on the Leica STELLARIS 8 confocal microscope using a 20x air objective (HC PL APO 20x/0.75 CS2) and a 63x oil-immersion objective (HC PL APO 63x/1.40-0.60 OIL). The data was visualized as maximum intensity projections using the Leica Las X software and ImageJ. The cells were quantified using ImageJ. The threshold was adjusted, holes filled, non-cells removed by using the binary open function and cells separated by the watershed function before quantifying the cells.

### IMpower150 trial sample RNA-seq analysis

To test whether pneumonitis incidence was correlated with tumor signatures at baseline prior to treatment, RNAseq transcriptomes from tumors biopsied prior to treatment within the IMPower150 trial were used in gene set enrichment analysis (GSEA). IMpower150 was an international, open-label, randomized, phase 3 trial conducted at 240 study centers in 26 countries (NCT02366143). The study was performed according to the Good Clinical Practice guidelines and the Declaration of Helsinki, with study protocol approval provided by independent ethics committees at each of the participating sites. Patients were randomly assigned, in a 1:1:1 ratio, to receive atezolizumab plus carboplatin plus paclitaxel (ACP group), atezolizumab plus bevacizumab plus carboplatin plus paclitaxel (ABCP group), or bevacizumab plus carboplatin plus paclitaxel (BCP group). Patient eligibility criteria and study design information have been published previously^38,39^. All patients provided written informed consent. Patients from the ACP arm in the IMPower150 trial (n=321) were stratified as pneumonitis positive versus negative for the GSEA analyses, and signatures were selected for a broad variety TME cell types^40^. GSEA was carried out using the qusage R package (v2.34.0)^59^ in R v4.3.1. Significant associations between pneumonitis incidence and tumor lesion RNA expression were then identified using the p-values (calculated using the probability density function for each gene set vs. the null hypothesis of no fold change as part of the qusage function) for each signature across the above/below median groups of patients. Race, sex and resection status were used as covariates in the GSEA, due to their known differential transcriptional differences.

### Single cell dissociation and single-cell RNA-sequencing (scRNA-seq) sample preparation

For the sequencing of the alveolar building block, TRMs, PBMCs and alveolar myeloid cells were thawed immediately before sequencing. Alveolar organoids were either grown in PneumaCult™ AvOE or PneumaCult™AvOD medium. The LIO and LIOM cultures were set up as described above and digested after 72 h of culture as published previously^25^. Briefly, organoids were removed from their matrigel domes and transferred to a 1% BSA-coated tube. The fragments were centrifuged and resuspended with enzymes from the Neural Tissue Dissociation Kit (Miltenyi Biotec). Every 5-7 min the suspension was pipetted with 1% BSA-coated tips to prevent cell loss. After 30 min the cells were filtered through a 40 µm Flowmi filter (Sigma-Aldrich) into another 1% BSA-coated tube. Next, the cells were centrifuged and resuspended in the magnetic microbeads of the Dead Cell Removal Kit (Miltenyi Biotec). For samples where epithelial cells had to be depleted, human EpCAM microbeads (Miltenyi Biotec) were added simultaneously. Dead cells and EpCAM+ cells were depleted on the MACS MS columns (Miltenyi Biotec) from the viable ones. The viable cells were filtered through a 70 µm Flowmi cell strainer, centrifuged and resuspended in an appropriate small volume of 1% BSA-HBSS buffer and counted on the Countess automated cell counter (invitrogen). The single cell libraries were prepared with the 10x Genomics platform using the Chromium Next GEM Single Cell 3ʹ Kit v3.1.

### Single-cell RNA-seq data preprocessing

CellRanger (v6.0.2, 10x Genomics) was used to extract unique molecular identifiers (UMIs), cell barcodes, and genomic reads from the sequencing results of 10x Chromium experiments. Then, count matrices, including both protein coding and non-coding transcripts, were constructed aligning against the annotated human reference genome (GRCh38, v3.0.0, 10x Genomics). In order to remove potentially damaged or unhealthy cells and improve data quality, the following filtering steps were performed in addition to the built-in CellRanger filtering pipeline. Cells associated with over 50,000 transcripts, cells associated with less then 500 unique transcripts detected together with cells characterized by over 20% of mitochondrial transcripts were removed. Transcripts mapping to ribosomal protein coding genes as well as mitochondrial genes were removed together with transcripts detected in less than 10 cells.

### Normalization with SCTransform

For normalization and variance stabilization of each scRNA-seq experiment’s molecular count data, we employed the modeling framework of SCTransform in Seurat v4^60^. In brief, a model of technical noise in scRNA-seq data is computed using ‘generalized gamma poisson regression’^61^. The residuals for this model are normalized values that indicate divergence from the expected number of observed UMIs for a gene in a cell given the gene’s average expression in the population and sequencing depth of the experiment. Additionally, a curated list of cell cycle associated genes, available within Seurat, was used to estimate the contribution of cell cycle and remove this source of biological variation from each dataset in order to increase the signal deriving from more interesting processes. The residuals for the top 2,000 variable genes were used directly as input to computing the top 100 Principal Components (PCs) by PCA dimensionality reduction through the RunPCA() function in Seurat. Corrected UMI, which are converted from Pearson residuals and represent expected counts if all cells were sequenced at the same depth, were log-transformed and used for visualization and differential expression (DE) analysis.

### Doublet removal with DoubletFinder

For each scRNA-seq experiment DoubletFinder^62^ was used to predict doublets in the sequencing data. In short, this tool generates artificial doublets from existing scRNA-seq data by merging randomly selected cells which are then pre-processed together with real data and jointly embedded on a PCA space that serves as basis to find each cell’s proportion of artificial k nearest neighbors (pANN). For this step we restricted the dimension space to the top 50 PCs. Finally, pANN values were rank ordered according to the expected number of doublets and optimal cutoff is selected through ROC analysis across pN-pK parameter sweeps for each scRNA-seq dataset; pN describes the proportion of generated artificial doublets while pK defines the PC neighborhood size. In order to achieve maximal doublet prediction accuracy, mean-variance normalized bimodality coefficient (BCmvn) was leveraged. This provides a ground-truth-agnostic metric that coincides with pK values that maximize AUC in the data. To overcome DoubletFinder’s limited sensitivity to homotypic doublets, we consider doublet number estimates based on Poisson statistics with homotypic doublet proportion adjustment assuming 1/50,000 doublet formation rate the 10x Chromium droplet microfluidic cell loading.

### Ambient mRNA signal removal

After doublet prediction and removal, we analyzed each scRNA-seq dataset in order to estimate the extent of ambient mRNA contamination in every single cell and correct it. We used the R package Cellular Latent Dirichlet Allocation (CELDA)^63^ which contains DecontX, a method based on Bayesian statistical framework to computationally estimate and remove RNA contamination in individual cells without empty droplet information. We applied the DecontX() function in CELDA to the raw count matrices with default parameters. Subsequently, we removed all cells with contamination values above 0.5 and we used the decontaminated count matrices resulting from DecontX() for downstream analysis.

### Geometric sketching

Geometric sketching is a downsampling technique that helps explore and interpret scRNA-seq data more effectively by providing a concise and intuitive representation of the cellular landscape that preserves rare populations. Individual datasets, after preprocessing, doublet removal and ambient mRNA regression, were aggregated according to specific criteria (individual building blocks, case/control of FOLR1 TCB, CytoStim treated conditions) and went through a joint normalization step with ScaleData() function in Seurat to mitigate technical confounding factors. This was followed by the selection of a set of meaningful 3,000 most variable global genes as implemented by FindVariableFeatures() in Seurat. We then used the sketchData() function from CellChat^64^, with default parameters, to select one third of the sequenced cells for each aggregated scRNA-seq dataset. This strategy was employed to avoid variability in sequence efficiencies that influence the computation of lower-dimensional embeddings and heterogeneity analysis.

### Joint processing of affine datasets

Selected cells after geometric sketching were jointly normalized using the SCTranform pipeline, described above, with 3,000 global variable genes selected and cell cycle signal removal. Normalized counts of global variable genes were subsequently used as input to computing the top 100 PCs through the RunPCA() function in Seurat. Leading 30 PCs and 50 nearest neighbors were then used to define the shared neighborhood graph with the FindNeighbors() function in Seurat. Subsequently, datasets were clustered according to the shared neighborhood graph using the Louvain algorithm^65^ through the Seurat function FindClusters() with resolution 0.2. Finally, top 50 PC vectors of the PCA space served as basis to obtain a two-dimensional (2D) representation of the data through Uniform Manifold Approximation and Projection (UMAP)^66^ implemented in RunUMAP() with 50 nearest neighbors. All identified clusters were manually characterized based on the expression of marker genes.

### Residency index

The residency index of tissue-resident as well as circulating T cells sequenced after isolation and prior to co-culture was computed by identifying the closest 50 neighbors of each T cell on the lower-dimensional space defined by the leading 50 PCA vectors. Subsequently, the proportion of PBMC-derived T cell neighbors was subtracted from the proportion of IIO-derived T cell neighbors of each individual cell and the resulting index was mapped to 0-1 interval.

### Differential expression analysis

Gene differential expression (DE) analysis between distinct cell populations in scRNA-seq data was assessed by performing Wilcoxon rank sum tests and auROC analysis as implemented by Presto package in R. Log-transformed corrected UMIs were used as input for the DE statistical tests, and genes were called differentially expressed if associated adjusted p-value (Bonferroni method) was lower than 0.05, AUC value was above 0.6 and log fold change was greater than 0.15. In addition, we also set thresholds on detection rates of DE genes. In particular, a given gene was assigned as over-expressed in the analyzed group if it was detected in at least 30% of the samples of that group, while the detection rate in the background samples was at most 70% of the detection rate of the analyzed group.

### Intercellular communication analysis

To investigate ligand-receptor (LR) mediated cell-cell communications during immune cell activation in response to different treatments, we extracted genes labeled as either ligands or receptors from curated databases^67^ and required that genes were differentially expressed between the various populations under investigation. This facilitated retrieval of directional information about the signal exchange. To gain insights into functional cell-cell communication, we used the NicheNet pipeline which considers the influence of sender-cell ligands on receiver-cell gene expression^67^.

### Functional enrichment analysis

To understand mechanisms underlying phenotypes in our data, differentially expressed genes were analyzed for gene ontology biological process (GOBP) enrichment using one-sided hypergeometric testing. P-values were adjusted for multiple testing hypotheses by the Bonferroni method and only enrichment results below a 5% significance level threshold were considered. For this analysis, we only considered biological processes consisting of sets with more than 10 but less than 300 mapped genes.

### Identification of drug responder groups

The Mixscape workflow provides a structured approach to identifying and analyzing the effects of perturbations at the single-cell level, helping to uncover the underlying biological mechanisms and responses. This model identifies the differential expression of genes in perturbed versus control cells and categorizes them into responder and non-responder groups based on their transcriptional profiles. In brief, we split the perturbation dataset into three cell populations including effector / memory T cells, immune-suppressive T regulatory cells, antigen presenting B and macrophage cells. Thus, we used CalcPertubSig() function from Seurat on the log-normalized counts with the leading 30 PC vectors (ndims = 30) and 20 nearest neighbors (num.neighbors = 20) to compute the perturbation signatures of each populations subject to different drug regimes compared to the CytoStim control. We then used the ScaleData() function from Seurat to normalize the computed perturbation signatures with do.scale and do.center set to FALSE and TRUE respectively, otherwise default arguments. Lastly, we invoked RunMixscape() function on the normalized perturbed signatures setting the de.assay to the SCTransform results, fine.mode argument to TRUE and fine.mode.labels argument to the cluster labels to identify the responder groups in each cell population and each drug treatment compared to the CytoStim control.

## Supplementary information

**Supplementary table**

The table reports information on antibodies used for all experiments.

**Supplementary video 1**

TRM cell migration within LIOs. Video generated from a confocal time-lapse experiment, depicting the movement of TRM cells (yellow) in co-culture with autologous human alveolar organoids (not rendered).

## Acknowledgements

We thank R. Sriram for supporting us with the establishment and oversight of the research agreements governing access to human lung specimens. We acknowledge the support of BeCytes for their continuous collaboration and support. We thank the Pathology Core Labs of pRED, Pathology and Applied Safety Science (PASS) for their support and technical guidance for the tissue based assays (IHC, mIF, whole slide scanning). We extend our gratitude to Mireia Serra Mitjants from the Thoracic Services of Hospital Universitario Mútua de Terrassa (Spain), the Donation and Transplantation Institute - DTI (Spain), and BeCytes Biotechnologies (Spain) for facilitating access to biospecimens and ensuring their availability for research in compliance with ethical and legal standards, following the provision of informed consent by the donor.

## Author contributions

L.S., L.C. and J.G.C. conceived the study. L.C., L.S., T.R., and J.G.C. wrote the manuscript. L.S., L.C., M.C.R. made the figures. L.S. was involved in designing, supervising and performing most experiments in the manuscript. L.S., M.P.M, M.B., L.G.T. performed scRNA-seq experiments. B.G. led the bioinformatics efforts and analysed all scRNA-seq data. Q.Y. supported the generation of visual graphs for the scRNAseq data. F.C. and L.K. performed the 3D phenotypic staining and image acquisition. B.Y. N. and V.G. analysed the clinical IMpower150 data. M.S. and A.D. exposed the myeloids to KLH and analysed the data. L.S. and E.L. performed the phagocytosis assays and proteome analysis of the myeloids. L.S. and G.L. performed cytokine measurements. L.S., R.O., J.N. stained cells and imaged them on confocal microscopes. L.S., M.P.M., L.L., T.Z., C.R., B.S. isolated and banked organoids, alveolar myeloids and TRM cells from primary lung samples. I.C., N.S-R, and C.Z. established and performed the IHC, mIF and HE stainings on human patient derived lung tissue and organoids. N.S-R. reviewed the quantitative image analysis performed by M.P.M and L.S. with HALO AI and evaluated the lung tissue morphology. N.G., A.B.R., R.M. supported the project with intellectual contributions.

## Declaration of generative AI and AI-assisted technologies in the writing process

During the preparation of this work, the authors used ChatGPT-4o to spell- and language-check written text. After using this tool, the authors reviewed and edited the content as needed and took full responsibility for the content of the publication.

## Availability of data and materials

All datasets supporting the conclusions of this article are included within the article (and its additional files).

## Code availability

Accession codes will be available before publication.

